# Bursting Translation on Single mRNAs in Live Cells

**DOI:** 10.1101/2022.11.07.515520

**Authors:** Nathan M. Livingston, Jiwoong Kwon, Oliver Valera, James A. Saba, Niladri K. Sinha, Pranav Reddy, Blake Nelson, Clara Wolfe, Taekjip Ha, Rachel Green, Jian Liu, Bin Wu

**Affiliations:** Department of Biophysics and Biophysical Chemistry, Johns Hopkins University School of Medicine, Baltimore, MD 21205, USA; Department of Molecular Biology and Genetics, Johns Hopkins University School of Medicine, Baltimore, MD 21205, USA; Department of Cell Biology, Johns Hopkins University School of Medicine, Baltimore, MD 21205, USA; The Center for Cell Dynamics, Johns Hopkins University School of Medicine, Baltimore, MD 21205, USA; The Solomon H Snyder Department of Neuroscience, Johns Hopkins University School of Medicine, Baltimore, MD 21205, USA; Howard Hughes Medical Institute, Chevy Chase, MD 20815, USA

**Keywords:** Single molecule, RNA, translation, stochastic gene expression, bursting

## Abstract

Stochasticity has emerged as a mechanism to control gene expression. Much of this so-called “noise” has been attributed to bursting transcription. However, the stochasticity of translation has not similarly been investigated due to a lack of enabling imaging technologies. We developed techniques to track single mRNAs and their translation in live cells for hours, allowing measurement of previously uncharacterized translation dynamics. We applied genetic and pharmacological perturbations to control translation kinetics. Like transcription, translation is not a constitutive process but instead cycles between inactive and active states or “bursts”. But unlike transcription, which is largely frequency modulated, complex structure in the 5’-untranslated region alters burst amplitude. Bursting frequency can be controlled through cap-proximal sequences and *trans*-acting factors such as eIF4F. We coupled single molecule imaging with stochastic modeling to deduce the fundamental kinetic parameters of translational bursting, a new dimension of translational control.

**Highlights:**

- Long-term tracking of single mRNAs reveals multi-state, bursting translation
- Structure in the 5’-untranslated region modulates translational burst amplitude
- 5’-cap proximal sequences modulate translational burst frequency
- mTOR signaling adjusts translation bursting to respond to environmental cues

## Introduction

Translation is the culmination of the central dogma of molecular biology where nucleic acid information is decoded into functional proteins. This process is highly regulated to deploy proteins when and where they are required. The chemistry of the translation process by the ribosome has been worked out in atomic detail (Ben-Shem et al., 2011; Jobe et al., 2019) and global measurements of ribosome footprints have provided an unprecedented view of the translational landscape (Ingolia et al., 2009; Ingolia et al., 2019). *In vitro* single molecule techniques using reconstituted bacterial and eukaryotic translation machineries have provided sensitive kinetic measurements of translation initiation (Tsai et al., 2012; Wang et al., 2019), elongation (Aitken and Puglisi, 2010; Blanchard et al., 2004a; Blanchard et al., 2004b), and termination (Lawson et al., 2021; Prabhakar et al., 2017), however direct observation of its temporal regulation in cells remains elusive. For example, for a given mRNA, is it constitutively translated at a given rate or dynamically adjusted in time? Does it spend more time translating or load more ribosomes per unit time? How do *cis*- and *trans*-factors influence these dynamics? How do cells integrate environmental signals to control translation of single mRNAs?

Translation initiation is a highly regulated, multistep process consisting of (i) RNA engagement by initiation factors and the small ribosomal subunit, (ii) ribosome scanning of the 5’-untranslated region (UTR) until recognition of a start codon, (iii) large ribosomal subunit joining and transition to the elongating phase (Sonenberg and Hinnebusch, 2009). For canonical, cap-dependent translation, the m^7^G cap is recognized by eukaryotic initiation factor 4E (eIF4E), which forms the eIF4F complex together with the DEAD box helicase eIF4A and the scaffolding protein eIF4G. The small ribosomal subunit is recruited by eIF4F and subsequently scans the 5’UTR until an AUG start codon (Gingras et al., 1999b; Hinnebusch, 2011). Elements in the

5’UTR such as RNA structure and upstream open reading frames act to regulate initiation efficiency and tune protein output of the main coding region (Hinnebusch et al., 2016; Kozak, 1986, 1989, 1991). The effect of these elements on *in vivo* translation kinetics involves probabilistic interactions between a suite of associated factors and is poorly understood.

Traditional ensemble methods average the properties of many cells and mRNAs and cannot distinguish the heterogeneous behavior of individual mRNAs. Gene expression results from a complex series of stochastic, single-molecule interactions leading to “bursty” transcription and noisy outcomes (Blake et al., 2003; Elowitz et al., 2002; Lenstra et al., 2016; Raser and O’Shea, 2004; Rodriguez et al., 2019; Sanchez and Golding, 2013; Suter et al., 2011; Zenklusen et al., 2008). Single cell mRNA levels appear to correlate with burst frequency (proportion of time transcription is active) as opposed to burst amplitude (number of mRNA produced per unit time) (Larson et al., 2013; Wan et al., 2021). For this reason, transcription is considered “frequency modulated”. Many of these insights have been garnered from single molecule RNA imaging, which accurately captures cell-to-cell and even allele-to-allele heterogeneity in temporal dynamics (Tutucci et al., 2018).

Our group and others have developed *in vivo* techniques to simultaneously track single mRNAs and their associated nascent peptides, providing real-time, single mRNA resolution of translation in living cells (Morisaki et al., 2016; Pichon et al., 2016; Wang et al., 2016; Wu et al., 2016; Yan et al., 2016). Single molecule imaging of nascent peptides (SINAPs) and similar techniques utilize an array of epitopes that, when translated, are co-translationally labeled by complementary single chain variable fragment fused to a mature fluorescent protein. The brightness of the nascent peptide (NAP) channel corresponds to the number of actively translating ribosomes. Stem-loop labeling systems are used to track single mRNAs in an independent channel. Co-translational labeling of NAPs on single mRNAs has been successfully employed to study diverse translational processes such as frameshifting (Lyon et al., 2019), translation-coupled mRNA decay (Hoek et al., 2019; Ruijtenberg et al., 2020), translational stress response (Mateju et al., 2020; Moon et al., 2019), non-canonical initiation (Boersma et al., 2019; Boersma et al., 2020; Koch et al., 2020; Wang et al., 2021b), and ribosome-associated quality control (Goldman et al., 2021). Despite much progress, little is known about the fundamental question of how single mRNAs are translated through time. This is because cytoplasmic mRNAs are highly mobile and cannot be tracked for the complete duration of translation events.

In this study, we employed long term tracking of single mRNAs and their associated translation sites to reveal the heterogeneity of translation in living cells. Like transcription, we observed translation is a discontinuous process that proceeds in “bursts” separated by translationally inactive “dwell” periods. Although translational bursting has been alluded to previously (Aguilera et al., 2019; Wu et al., 2016), it has yet to be quantified and mechanistically dissected. We designed reporters to strategically perturb those bursting behaviors. First, we fixed the 5’UTR but changed the length of open reading frame (ORF) to firmly establish that a single constant initiation process is unable to recapitulate the presence of translational bursts, and requires we invoke a multi-state initiation model to explain the presence of translationally inactive periods. Next, we added defined structures to the 5’UTR to establish that structure alters protein output by modulating burst amplitude as opposed to timing. Finally, we determine that cap-proximal sequences control burst frequency and response to environmental stimuli. We coupled our experiments with stochastic modeling to determine the fundamental rates of translation initiation and bursting. Taken together, our study adds a new dimension to our understanding of translation temporal dynamics in its native context.

## Results

### Long term imaging of single mRNAs and translation sites

To interrogate the long-term translational activity of single mRNAs *in vivo*, we employed a modified version of the SINAPs technology developed in our group and others (Morisaki et al., 2016; Pichon et al., 2016; Wang et al., 2016; Wu et al., 2016; Yan et al., 2016). The SINAPs reporter mRNA contains tandem repeats of SunTag (GCN4) epitopes in the N-terminus of the ORF that, when translated, are bound outside of the ribosomal exit tunnel by a stably expressed single-chain variable fragment antibody against GCN4 fused to a super-folder GFP (scFv-sfGFP) (Tanenbaum et al., 2014). Downstream of the SunTag repeats, we added luciferase or fluorescent proteins to facilitate ensemble gene expression measurements. The C-terminus of the SINAPs ORF contains an auxin inducible degron (AID) that is selectively marked for degradation in the presence of indole acetic acid (IAA) by the stably expressed OsTIR1 E3 ubiquitin ligase (Nishimura et al., 2009; Wu et al., 2016). Depletion of released mature proteins allows for recycling of scFv-sfGFP for labeling and reducing background fluorescence, essential for detecting single peptides to calibrate the number of nascent peptides at translation sites in fixed cell measurements (Latallo et al., 2019).

To visualize single mRNAs in live cells, the 3’UTR of the SINAPs reporter contains an array of bacteriophage-derived stem loops, usually non-repetitive, sequence-synonymized MS2 or PP7 binding sites (MBS, PBS) (Wu et al., 2015). These stem loops are bound by a complementary MS2 coat protein (MCP) or PP7 coat protein (PCP) fused to a fluorescent protein (Wu et al., 2012). The engineered dihaloalkane dehalogenase “HaloTag” is widely used in single molecule imaging owing to its ability to conjugate the bright, highly photostable Janelia Fluor dyes and is fused to either MCP or PCP for mRNA imaging (Grimm et al., 2016; Grimm et al., 2021; Los et al., 2008).

A single round of translation takes place on the time scale of minutes, and multiple cycles may take tens of minutes. Despite improvements to imaging technologies, tracking freely diffusing single mRNAs for multiple rounds of translation remains a significant challenge. Tethering mRNA to the plasma membrane using anchored MCP or PCP molecules extended the observation window without influencing translation efficiency and protein output (Goldman et al., 2021; Yan et al., 2016) but still suffered from limited track length, ambiguous identification of non-translating mRNAs from aggregating membrane coat proteins, and high background signal due to unbound coat proteins on the membrane. To address these issues, we separated mRNA fluorescent tagging from membrane tethering components (Figure 1A). Our modified 3’UTR contained an array of 24xMBS followed by 12xPBS. The MBS were labeled co-transcriptionally in the nucleus by a tandem dimeric MCP-HaloTag fusion protein (tdMCP-HaloTag) and the PBS were anchored to the plasma membrane by binding to unlabeled tdPCP-SNAPtag-CAAX (Figure 1A) (Wu et al., 2012). We label tdMCP-HaloTag with JFX-549 to illuminate the RNA in the red channel (Grimm et al., 2021). Compared with direct tethering with MBS, we observed ∼2x increase in signal-to-background (Figure 1B and Figure S1A) and increased mean RNA tracking times (from 48.6 to 67.4 minutes) when considering tracks longer than 30 minutes (Figure 1C and Figure S1B). More tracks (22%) exceeded one hour in the MBS-PBS system compared with that of MBS alone (<4%). Using a SINAPs reporter coding for a minimal ORF of 24xSunTag-AID (ST-AID), we achieved a mean translation site tracking time of 83.2 minutes.

**Figure 1.**
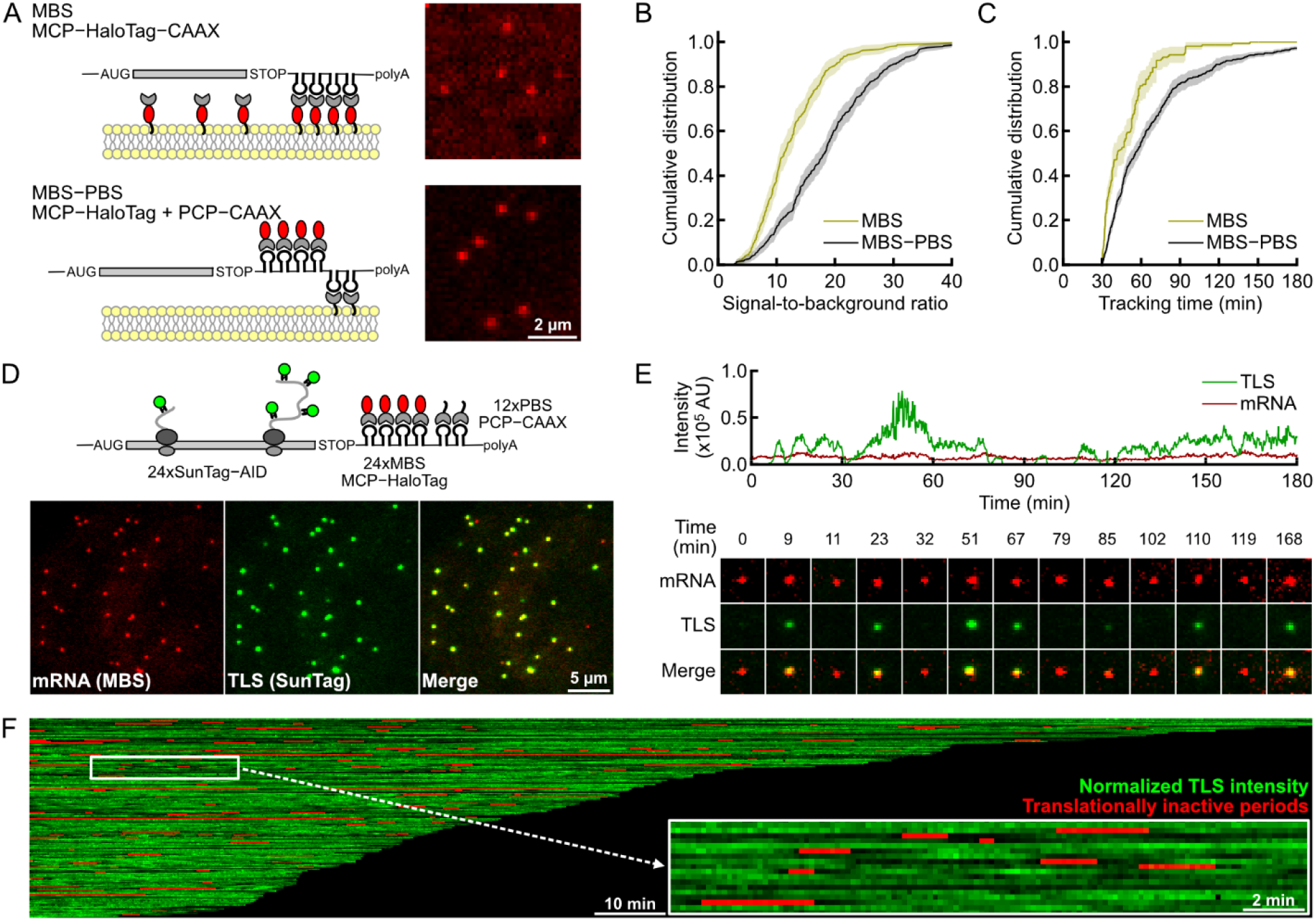
Translational bursting observed with long term single mRNA tracking. (A) Membrane tethering of translating mRNAs using coat proteins targeted to plasma membrane. Top: direct MBS tethering. Bottom: MBS-PBS dual stem-loops tethering. Right: example images. (B) MBS-PBS dual tethering increased signal-to-background of mRNA signals compared with MBS direct tethering (n=247-374 mRNA, CDF ± 95% CI). (C) MBS-PBS tethering achieves longer tracking time than direct MBS tethering (n=150-439 mRNAs, single cell mean ± SEM). (D) Top: scheme for the ST-AID reporter. Bottom: representative images for membrane tethered mRNAs (Red) and translation sites (TLS, Green). (E) Top: sample TLS trajectory exhibits on-to-off transitions. Bottom: images of the mRNA (Red) and its TLS (Green). (F) Composite of single mRNA translation trajectories for the ST-AID reporter (n=168 mRNA). Insert: zoomed view of single mRNA tracks. CDF: cumulative distribution function; CI: confidence interval; SEM: standard error of the mean.

With increased tracking time, we observed that translation is a highly dynamic and heterogenous process characterized by active and inactive periods (Figure 1D-F and Video S1). This observation was consistent with previous studies using SINAPs in cultured primary neurons where relatively stationary dendritic mRNAs fluctuated between active and inactive translation on time scales in the tens of minutes (Wu et al., 2016). Pre-treating cells with the translation elongation inhibitor cycloheximide prior to imaging abolishes bursting behavior (Figure S1C, D).

We extracted three experimental parameters from our measurements: burst duration. dwell duration, and burst amplitude. Translation site intensity traces of single mRNAs were segmented into active bursts and inactive dwells (Figure 2A). This binary analysis identified the timing of translational bursts (Figure 2B, C and STAR Methods). The measured mean durations of bursts and dwells are 43.3 min and 3.5 min, respectively, for the ST-AID reporter (Table S1).

**Figure 2.**
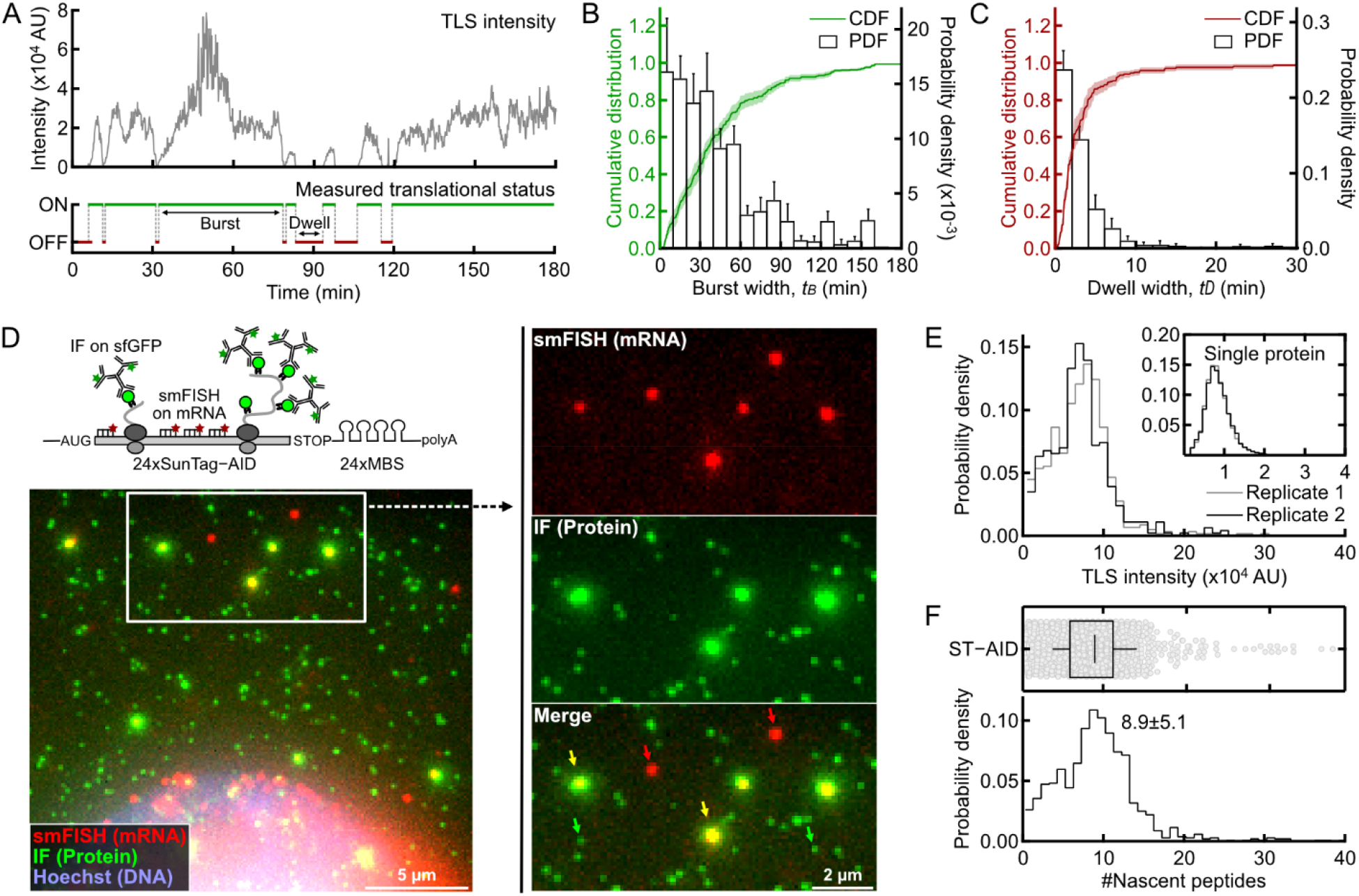
Measuring burst parameters. (A) Segmentation of TLS intensity traces into active (burst) and inactive (dwell) periods. (B-C) The cumulative and probability distributions functions of burst (B) and dwell (C) periods (n=168 mRNAs, single cell mean ± SEM for both displays). (D) Fixed cell measurement of single mRNAs, TLS and released single polypeptides using smFISH-IF. Green: protein. Red: mRNA. Blue: DNA. Enlarged box represents indicated region. The green, red and yellow arrows in the right panel indicate mature single proteins, non-translating mRNAs and TLS, respectively. (E) The distribution of integrated intensities of TLS and single proteins (insert) of two biological replicates of smFISH-IF experiments expressed in arbitrary units prior to normalization (n=1,047 TLS, 41,228 single proteins). (F) The TLS intensities are normalized using released polypeptides from the same cell to calculate the number of nascent peptides. Top: beeswarm plot. Bottom: histogram distribution (box plot represents median ± interquartile range, whisker for SD). AU: arbitrary unit; SEM: standard error of the mean.

To measure burst amplitude or ribosome load, we used single molecule fluorescence *in situ* hybridization coupled with immunofluorescence (smFISH-IF) to identify the distribution of translation site intensities in units of nascent peptides (Latallo et al., 2019). The mRNA was labeled with fluorescent probes targeting the SunTag region of the reporter, while scFv-sfGFP was labeled with an anti-GFP antibody (Figure 2D). Discrete fluorescent spots in protein channel not colocalized with RNA spots represented single mature proteins (Figure 2D, green arrows). The bright puncta colocalized with mRNAs were translation sites (TLS), whose integrated intensities were normalized by the single proteins in the same cell to measure the number of NAPs (Figure 2E, F and STAR Methods). This measurement is highly reproducible from day-to-day, with a mean of 8.9 NAPs for the ST-AID reporter (Figure 2E, F and Table S1). Addition of 12xPBS and membrane targeting did not affect the number of nascent peptides associated with the mRNA (Figure S2B). Ribosomes actively translating the SunTag region may have only partially synthesized the SunTag array, leading to an underestimation of ribosome number based on the normalized TLS intensity (STAR Methods and Figure S2C); with this consideration, we can correct the NAP measurement based on theoretical models (STAR Methods). The reported NAP value is uncorrected, because it is model-independent.

### A two-state model of translational bursting

When the average number of ribosomes on an mRNA is low, it is anticipated that a given mRNA may have stochastic excursions into a state without any ribosomes. To examine whether the measured translation dynamics can be described by a constitutive initiation process (where the initiation is random but the rate is constant), we performed stochastic Monte Carlo simulation based on the totally asymmetric exclusion process (TASEP) (STAR Methods and Figure S3A-C) (Zia et al., 2011). Given the average number of NAPs on mRNAs, we estimated the probability for the mRNA to reach zero ribosomes (STAR Methods). For the ST-AID reporter (the estimated initiation rate is = 4.4 min^-1^, corresponding to a mean NAPs density of 8.9), the mRNA has extremely low probability (<0.001) of transiting to the zero-ribosome state (Figure S3C) under a constitutive initiation model. Thus, we conclude that the observed dwell periods cannot be explained by a simple constitutive initiation process.

Next, we adopted the simplest multi-state model: a random telegraph model, that has been extensively used in the transcription field (Raj et al., 2006; Zenklusen et al., 2008; Zenklusen and Singer, 2011) (Figure 3A). In this model, an mRNA can exist in two intrinsic states, bursting (active) and dwelling (inactive). Only in the active state the mRNA can be translated. However, the observed fluorescence translation signal may deviate from the intrinsic state (STAR Methods and Figure S3D-F). For example, in a short active state, there may be no translation event if the initiation rate is low which results in a longer observed dwell period. On the other hand, if a short inactive state is between two active states, the mRNA may show a merged, longer measured burst period because the ribosomes in the first active state may not have finished elongation by the time the subsequent burst begins (i.e., overlap between bursts) (Figure 3A).

**Figure 3.**
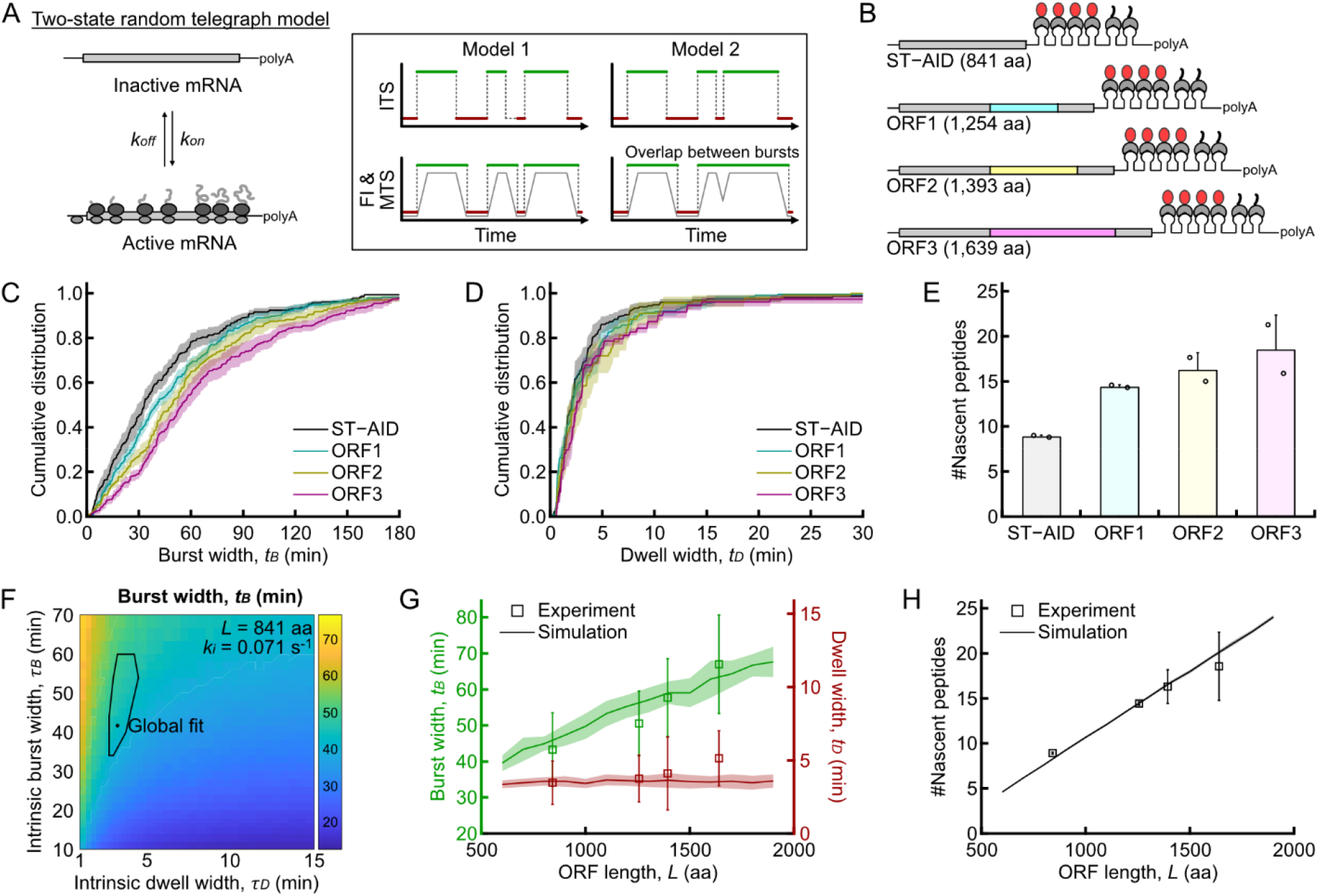
A two-state model for translational bursting. (A) Left: mRNAs transition between translationally active and inactive states. Model 1, no overlap between bursts and new translation burst only initiate on empty mRNAs. In this case, the measured burst parameter is similar to the intrinsic parameter (Figure S3D). Model 2, overlap is allowed and stochastic switching between states is independent of the number of ribosomes on the mRNA. In this case, intrinsic burst status may deviate from measured burst status depending on model (Figure S3E). (B) Reporters with different ORF lengths, but identical 5’ and 3’UTRs. (C) Distributions of measured burst times exhibits strong ORF length dependence (n=168-235 mRNAs, single cell mean ± SEM, ***p*** = 0.0014). (D) Distributions of measured dwell time are similar between different ORF lengths (***p*** > 0.05). (E) Average number of NAPs increases with ORF length (n=419-1,291 TLS, mean of 2 biological replicates ± SD, ***p*** = 0.0396). (F-H) Global fit of 4 reporters using phase diagram analysis uncovers intrinsic burst and dwell parameters. Error region (black line) represents confidence intervals (see STAR Methods). A single set of parameters of Model 2 (overlap allowed) described the ORF length dependent of observed mean burst and dwell times (G, mean ± SD), and number of NAPs (H, mean ± SD). SEM: standard error of the mean; SD: standard deviation. ***p*** values were obtained from one-way ANOVA test.

We considered two different cases for this off-to-on transition. First, bursts do not overlap, and the active state only begins with an empty mRNA (Figure 3A, Model 1). Second, bursts can overlap, and the state transition is independent of ribosome loading, meaning that the mRNA can be turned on even when elongating ribosomes from the previous burst are still present (Figure 3A, Model 2).

To distinguish these two models, we generated a series of constructs with different ORF lengths by inserting protein motifs between the SunTag and AID in the SINAPs reporter. The final lengths of ORFs including the linkers are 841, 1254, 1393, and 1639 amino acids for ST-AID, ORF1, ORF2, and ORF3, respectively. In Model 1, measured burst and dwell times are independent of ORF length (Figure S3D). However, Model 2 predicts that longer ORFs produce longer measured burst times because there is a higher probability of overlap between bursts (Figure S3E). In both models, the dwell is insensitive to the ORF length (Figure S3D-F). Experimentally, the measured burst widths were clearly extended for the longer ORFs whereas the measured dwell widths showed no significant differences (Figure 3C, D, Figure S3G, H and Table S1), agreeing with the prediction of Model 2. The measured average number of NAPs by smFISH-IF also increased with ORF length (Figure 3E, Figure S3I-L and Table S1). Consequently, Model 2 agrees with our data where bursts can overlap and the measured burst width may be longer than the intrinsic on period of the mRNA (STAR Methods). Because the measured parameters of the shortest ST-AID reporter most closely mirror the intrinsic bursting characteristics, we proceeded to use this ORF in all following experiments unless explicitly stated otherwise.

To extract the three intrinsic kinetic parameters for the random telegraph model of translation, intrinsic burst time (*τ*_*B*_), dwell time (*τ*_*D*_) and the initiation rate (*k*_*i*_), from the experimental data, we performed Monte Carlo simulation to generate a phase diagram of measured burst time (*t*_*B*_), dwell time (*t*_*D*_) and the average number of NAPs over the potential parameter space (*k*_*i*_, *τ*_*B*_ and *τ*_*D*_; for the four different ORF lengths) (Figure 3F). For example, the measured burst width depends on both intrinsic burst time and intrinsic dwell. For a given intrinsic burst time, the measured burst time increases when the intrinsic dwell time decreases as there will be more overlapping events. We compared the experimental results (*t*_*B*_, *t*_*D*_ and the number of NAPs) with the simulated data to extract the intrinsic parameters (*τ*_*B*_, *τ*_*D*_ and *k*_*i*_).

We performed a global fit of all measured burst times, dwell times and the number of NAPs for the four constructs with the same 5’ and 3’UTRs but different ORF lengths, assuming they have the same intrinsic parameters (*τ*_*B*_, *τ*_*D*_ and *k*_*i*_). A single set of intrinsic parameters (*k*_*i*_ = 4.3 min^-1^, *τ*_*B*_ = 42 min and *τ*_*D*_ = 3.3 min; Table S1) can describe all the measured quantities (Figure 3F-H). Specifically, the measured burst time (*t*_*B*_) and the average number of NAPs increase with the ORF length, as predicted by the two-state random telegraph model (Figure 3G, H).

### Structure in the 5’UTR modulates burst amplitude

The 5’UTR plays an important role in determining the translational efficiency of the downstream ORF (Hinnebusch et al., 2016). Secondary structure in the 5’UTR is thought to hinder 43S scanning and reduce initiation efficiency. We hypothesized that 5’UTR RNA structures could modulate translation bursting parameters and therefore protein output by two mechanisms (Figure 4A). In Model A, 5’UTR structure modulates burst frequency; a stronger RNA structure causes a longer dwell period because it takes longer to dissolve. In this first case, relative protein output could be explained through changes in burst timing. In Model B, 5’UTR structure modulates burst amplitude; a stronger RNA structure controls the number of ribosomes loaded per unit time and it takes longer for a 43S to scan through and initiate. In the second case, relative protein output could be explained by changes in the burst amplitude.

**Figure 4.**
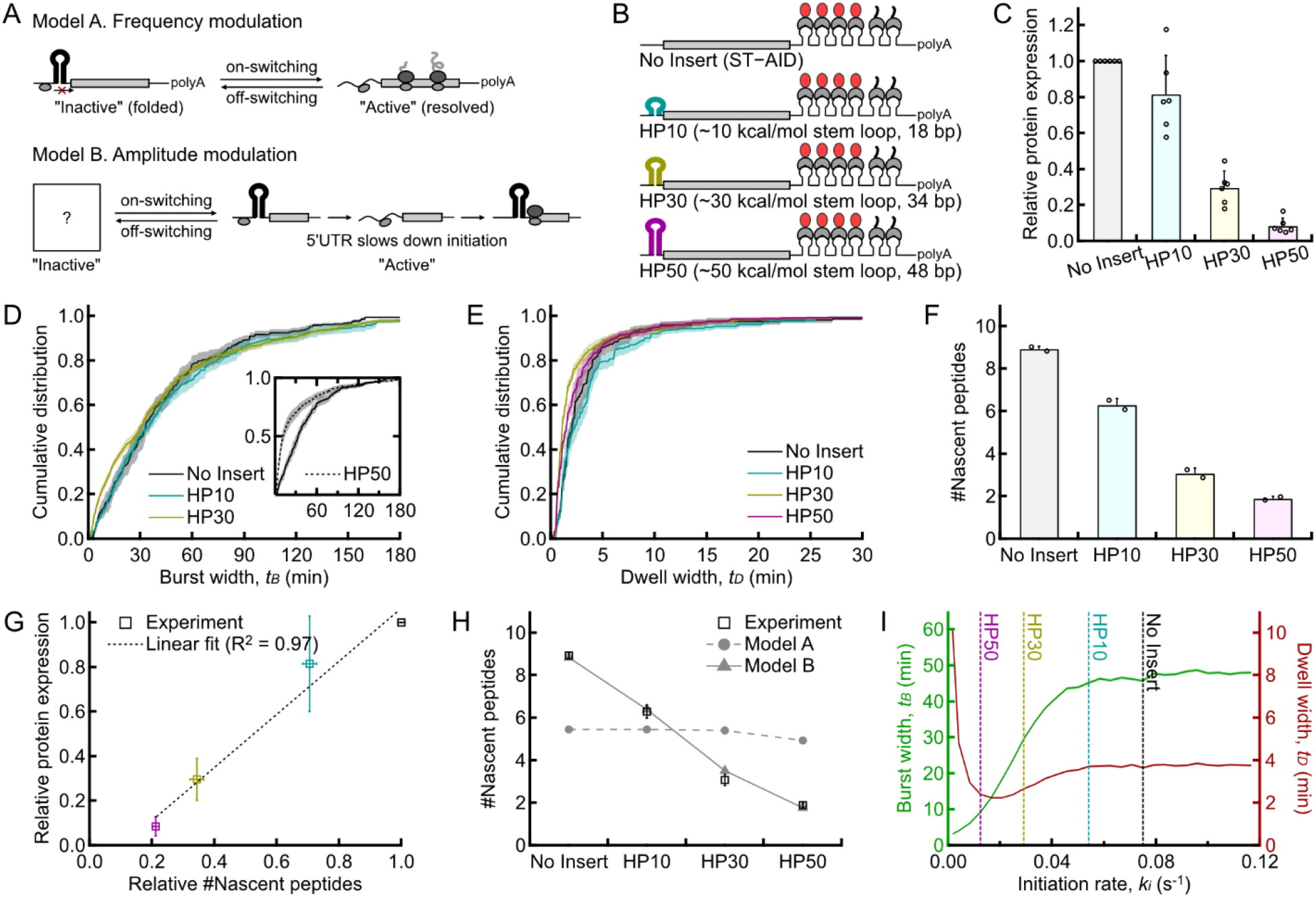
5’UTR hairpin secondary structures control burst amplitude. (A) Two competing models of how secondary structure in 5’UTR reduces protein output: either changing burst frequency (Model A) or burst amplitude (Model B). (B) Hairpins with different thermal stability were inserted in the 5’UTR. The reporters have the same ORF and 3’UTR. (C) Protein output, measured by dual luciferase normalized by RNA levels, decreased with the strength of hairpin (mean of 6 biological replicates ± SD, ***p*** < 0.0001). (D-E) 5’UTR secondary structure has a minimal effect on burst and dwell times (n=168-643 mRNAs, single cell mean ± SEM, ***p*** > 0.05 for both burst and dwell widths excluding the burst data of HP50). HP50 burst time is reduced due to low initiation rate (D, insert), as explained in panel I. (F) Hairpin strength dependent reduction in the number of NAPs (n=493-1,509 TLS, mean of 2 biological replicates ± SD, ***p*** > 0.05 for both burst and dwell widths excluding the burst data of HP50). (G) Total protein output correlates linearly with the number of NAPs, data from panel C and F. (H) The number of NAPs (symbol, experimental data from panel F) agrees with Model B (solid line), where hairpins control burst amplitude. Model A (dashed line) cannot explain the observed number of NAPs. (I) Monte Carlo simulation of burst and dwell time at different initiation rates in the active state. At high initiation rate, the burst and dwell times reach plateau. At low initiation rates, there are more zero ribosome events by chance, leading to shorter observed burst periods. SEM: standard error of the mean; SD: standard deviation. ***p*** values were obtained from one-way ANOVA test.

To systematically test the effect of secondary structure on bursting parameters, we inserted hairpins of increasing thermal stabilities in the 5’UTR immediately before the Kozak sequence (Babendure et al., 2006) (Figure 4B). We named these constructs HP10, HP30 and HP50 corresponding to their predicted thermal stability in kcal/mol. Using SINAPs constructs containing a Nanoluciferase insert, we performed luciferase and RT-qPCR to determine the effect of the hairpins on protein output (STAR Methods). As expected, the protein level was inversely related to hairpin strength (Figure 4C).

We performed both live cell and fixed cell imaging to measure translational bursting parameters. Surprisingly, constructs with different hairpins have very similar burst and dwell time distributions (Figure 4D, E, Figure S4A, B and Table S1) apart from the strongest hairpin HP50 which showed a modestly reduced burst width (Figure 4D, inset). In contrast, we observed the number of NAPs on the mRNAs decreased significantly with increased hairpin strength (HP10, HP30 and HP50 decreased 29%, 66% and 79% respectively compared with No Insert) (Figure 4F, Figure S4C-F and Table S1), and protein production is linearly correlated with the burst amplitude (Figure 4G).

To definitively distinguish the amplitude and frequency modulation models (Figure 4A), we performed global fit analysis for all constructs with different 5’UTR hairpins using the simulated phase diagram. In Model A, all hairpin constructs shared the same initiation rate but had different intrinsic burst and dwell widths. In Model B, all constructs shared intrinsic burst and dwell widths, but had distinct initiation rates depending on hairpin strength. Model A fails to recapitulate the ribosome load even though the intrinsic burst and dwell widths were tuned to fit each construct (Figure 4H, dashed line). In Model B, the initiation rate is adjusted to match the measured number of NAPs (Figure 4H, solid line) and a single set of intrinsic burst and dwell times are used to recapitulate the measured burst and dwell time. Importantly, when the initiation rate is low (<0.05 s^-1^), the measured burst time is reduced (Figure 4D, inset). This is because when the average number of ribosomes is low, mRNAs may occasionally have zero ribosomes, resulting in the reduction of both measured burst and dwell widths (STAR Methods), as recapitulated by the Monte Carlo simulation (Figure 4I). These stochastic zero-ribosome events break the intrinsic burst into smaller measured burst fragments. Taken together, these data suggest that the structures in 5’UTR change burst amplitude, rather than burst frequency.

### 5’TOP RNA sequences modulate burst frequency

TOP mRNAs contain a 5’ Terminal Oligopyrimidine tract (5’TOP) and encode translation factors and ribosomal proteins. These mRNAs are acutely responsive to nutrient and growth dependent translational control (Meyuhas and Kahan, 2015). Mechanistic target of rapamycin (mTOR) is the catalytic subunit of both the mTORC1 and mTORC2 complexes in mammalian cells and is a master regulator of cell growth and response to environmental changes. mTORC1 is responsible for regulating translation at multiple nodes (Hay and Sonenberg, 2004; Ma and Blenis, 2009). Active mTORC1 directly phosphorylates eIF4E binding proteins (4E-BPs), preventing their inhibitory association with eIF4E, and allowing formation of active eIF4F complex on the mRNA cap to begin the multi-step process of translation initiation (Gingras et al., 1999a; Pause et al., 1994; Sonenberg et al., 1978).

TOP mRNA translation is acutely affected by mTOR signaling (Hsieh et al., 2012; Jefferies et al., 1994; Meyuhas and Kahan, 2015; Thoreen et al., 2012; Yamashita et al., 2008). Interestingly, TOP mRNAs exhibit an “all-or-nothing” translation phenotype. Under periods of low nutrient availability these mRNAs exhibit a bimodal distribution between sub-polysome and polysome fractions (Hornstein et al., 2001; Meyuhas et al., 1987). We hypothesized that this phenotype would correlate with the bursting translation that we observe here and confer unique bursting characteristics when compared with our previous, non-TOP 5’UTR.

We generated a new construct containing the core-promoter and the 5’UTR of eukaryotic elongation factor 1A (EEF1A), a known TOP mRNA (Figure 5A). The translation intensity traces of EEF1A showed more bursting compared to our No Insert reporter, defined by longer inactive (red) and shorter active periods (green) (Figure 5B, C and Video S2). The mean burst and dwell widths of EEF1A were 31 min and 5.7 min, respectively (Figure 5C, D, Figure S5A, B and Table S1). The average ribosome loads measured by smFISH-IF, on the other hand, are similar to those of our control No Insert reporter (9.5 and 9.3 mean number of NAPs for No Insert and EEF1A, respectively) (Figure 5F, Figure S5C-E and Table S1).

**Figure 5.**
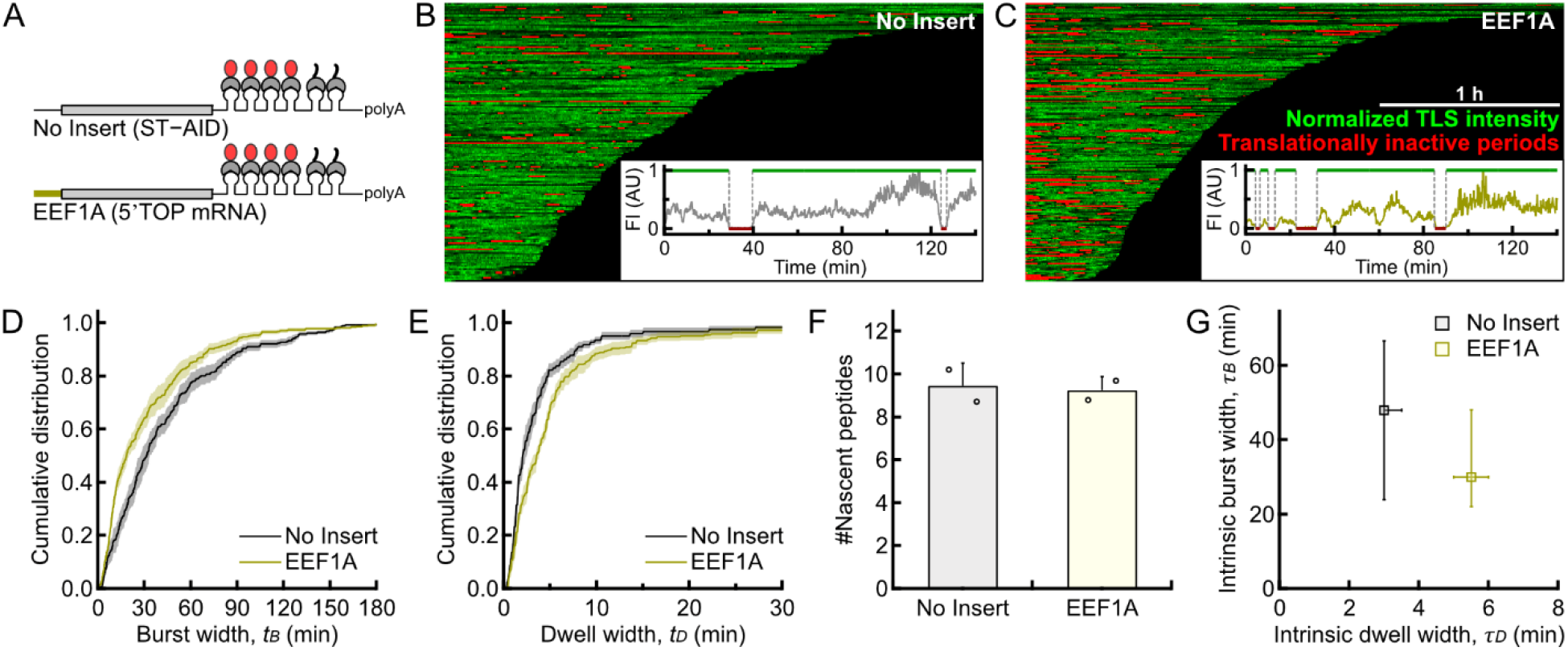
TOP mRNA exhibits unique burst transitions. (A) A reporter TOP mRNA with the EEF1A core promoter and 5’UTR but the same ORF and 3’UTR as the No Insert reporter. (B-C) Bursting heatmaps and sample traces (insert) of No Insert and EEF1A reporters (n=168-209 mRNA). (D-E) EEF1A exhibits shorter observed burst times and longer observed dwell times than the No Insert reporter (n=168-545 mRNAs, single cell mean ± SEM, ***p*** = 0.0270 and 0.0280 for burst and dwell widths, respectively). (F) The number of NAPs of No Insert and EEF1A are similar (n=1,037-1,205 TLS, mean of 2 biological replicates ± SD, ***p*** > 0.05). (G) Intrinsic burst parameters for No Insert and EEF1A extracted from phase diagram analysis. Error bars represent confidence intervals (see STAR Methods). SEM: standard error of the mean; SD: standard deviation; ***p*** values were obtained from Student’s t test.

The intrinsic parameters extracted from the phase diagram analysis revealed burst time *τ*_*B*_ = 30 min and dwell time *τ*_*D*_ = 5.5 min for EEF1A, clearly distinct from the No Insert reporter (Figure 5G and Table S1) but maintaining a similar amplitude (Fig. 5F). Contrasting with our amplitude modulation model for 5’UTR structure, we found that TOP sequences alter only the temporal dynamics of translational bursting.

### mTOR inhibition modulates both bursting frequency and amplitude

To further explore the role 5’UTRs in bursting, we inhibited mTOR signaling using Torin-1 to probe the effect of this signaling axis on bursting parameters (Figure 6A). Torin-1 is an ATP-competitive mTOR inhibitor with a cellular IC50 in the low nanomolar range (Liu et al., 2010). Under 250 nM Torin-1 treatment, the mTORC1 complex phosphorylated 4E-BP levels were reduced within 30 minutes and plateaued between 2-5 hours, while the reduction of phosphorylated S6K reaches maximal response within 15 minutes (Figure S6A, B). We pretreated cells with Torin-1 and imaged between 2-5 hours following treatment.

**Figure 6.**
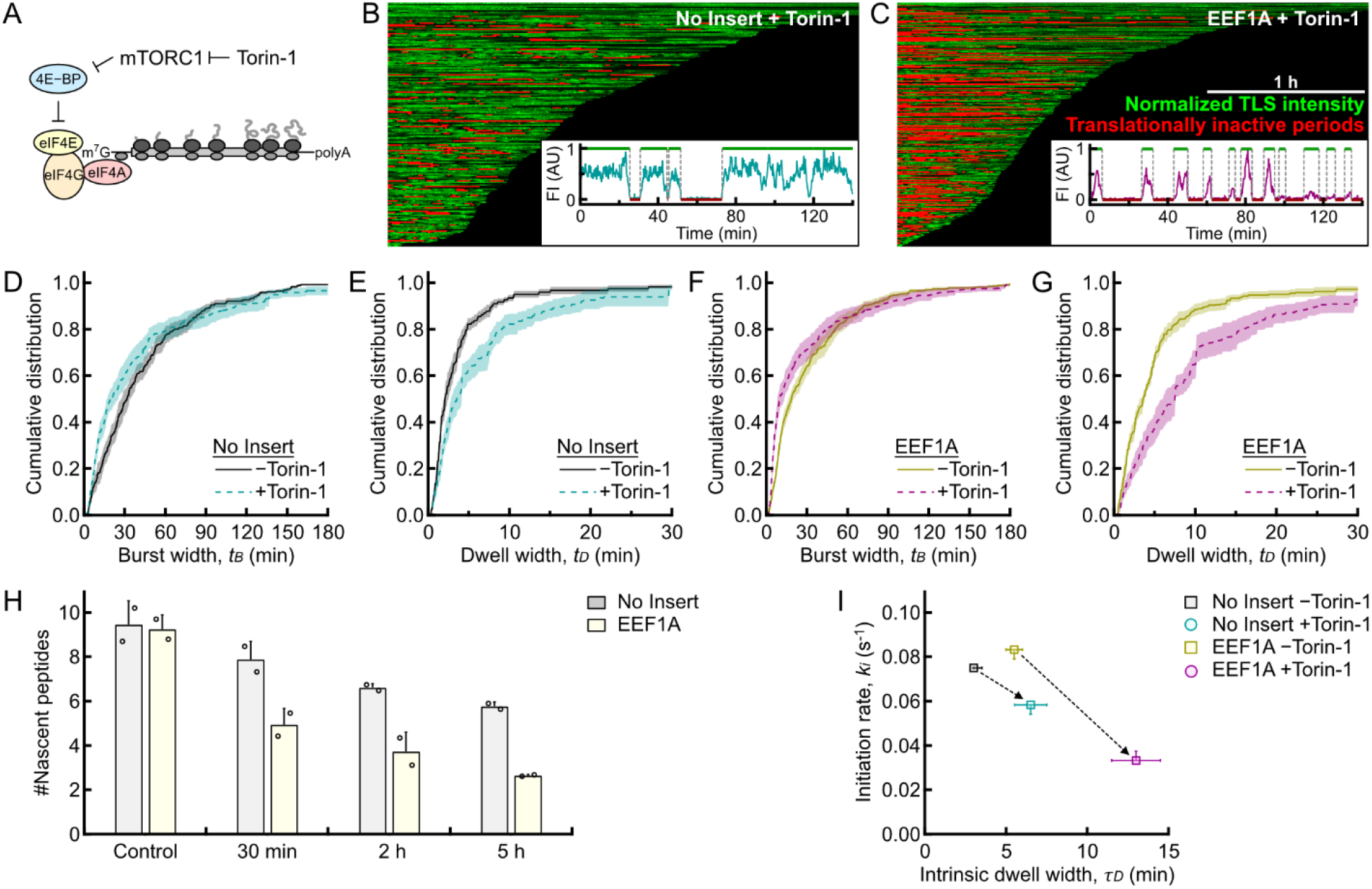
Torin-1 inhibition of mTOR signaling alters burst timing and amplitude. (A) Torin-1 inhibits the kinase activity of mTORC1, inhibiting translation initiation. (B-C) Heatmap analysis and sample TLS intensity traces (insert) of No Insert (B) and EEF1A (C) (D-G) Torin-1 treatment reduced observed burst times, and significantly increases dwell times for No Insert (n=168-184 mRNA, ***p*** > 0.05 for burst widths and ***p*** = 0.0164 for dwell widths) and EEF1A (n=296-410 mRNA, single cell mean ± SEM, ***p*** > 0.05 for burst widths and ***p*** = 0.0062 for dwell widths). (H) Number of NAPs measured at time points following treatment with Torin-1 or DMSO (n=85-1,812 TLS, mean of 2 biological replicates ± SD, ***p*** = 0.0206 and 0.0020 for No Insert and EEF1A, respectively). (I) Initiation rate and intrinsic dwell time for No Insert and EEF1A following Torin-1 treatment, extracted from phase diagram analysis. Error bars represent confidence intervals (see STAR Methods). SEM: standard error of the mean; SD: standard deviation. ***p*** values were obtained from Student’s t test (D-G) or one-way ANOVA test (H).

Torin-1 induced more periods of translational inactivity for both No Insert and EEF1A reporters, with EEF1A mRNA transitioning more frequently than No Insert (Figure 6B, C and Video S3-4). Following Torin-1 treatment, measured burst times for both constructs were reduced (No Insert: 43 → 39 min; EEF1A: 31 → 28 min), whereas dwell times were both significantly increased (No-Insert: 3.5 → 6.8 min; EEF1A: 5.7 → 10.6 min) (Figure 6D-G, Figure S6D, E and Table S1). Inhibition of mTORC1 signaling also altered burst amplitudes for both constructs (Figure 6H, Figure S6F-L and Table S1), but EEF1A responded faster and to a greater extent, consistent with the notion that TOP mRNAs are more sensitive to mTOR inhibition by Torin-1 (Hsieh et al., 2012; Thoreen et al., 2012).

Inhibiting mTOR by Torin-1 decreases the concentration of active eIF4F complex. We hypothesize that the translationally active state is when the mRNA is bound by active eIF4F. If the concentration of active eIF4F complex is reduced upon Torin-1 treatment, the inactive mRNA needs to wait longer to bind to eIF4F. However, once eIF4F is formed on the cap, its dissociation may be independent of Torin-1 treatment. Based on this potential mechanism, we hypothesized that mTOR inhibition by Torin-1 would only control the intrinsic dwell time and the initiation rate but would not affect the intrinsic burst time. We extracted the intrinsic bursting parameters using the phase diagram analysis, and the resulting measured values agreed well with the experimental observations (Figure S6M, N). Torin-1 increased the intrinsic dwell widths (No Insert: 3.0 → 6.5 min; EEF1A: 5.5 → 13 min) and reduced the initiation rates (No Insert: 4.5 → 3.5 min^-1^; EEF1A: 5.0 → 2.0 min^-1^) (Figure 6I and Table S1). The slight reduction in the measured burst widths is a result of the increased dwell widths, thus reducing the probability of overlaps and was well explained by Monte Carlo simulations without perturbing the intrinsic burst widths.

## Discussion

Initiation is often cited as the rate limiting step of translation, though the temporal dynamics of this process have not yet been thoroughly investigated in real-time inside cells. We advanced our SINAPs technology to directly observe multiple rounds of translation on single mRNAs. We coupled these experimental measurements with stochastic modeling to elucidate the fundamental parameters of translational initiation. We found that translation is a discontinuous process that fluctuates between translationally active and inactive states, requiring that we invoke a multi-state initiation model (Figure 7). This multi-state model is analogous to transcription, where factors such as chromatin state, transcription factor binding and enhancer-promoter interactions dictate bursting parameters (Rodriguez and Larson, 2020). Using measurements of translational burst time, dwell time and burst amplitude, we set out to understand how these parameters can change in response to known mechanisms of translational initiation control.

**Figure 7.**
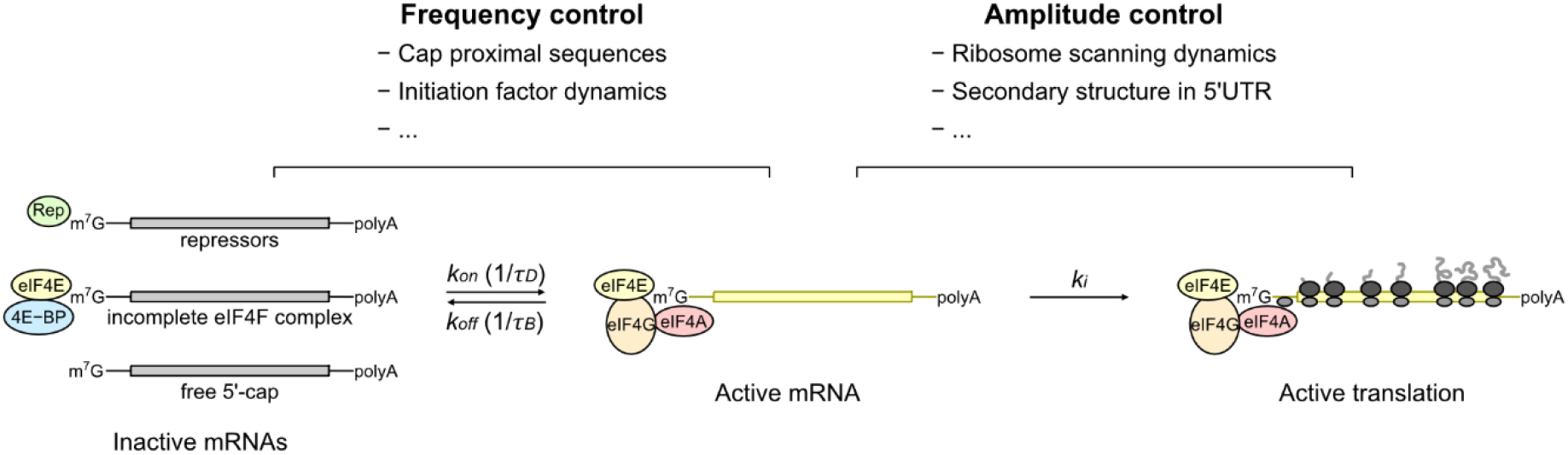
A working two-state model of translation initiation. An mRNA transitions between active and inactive states. Once activated, either by binding of activator complex or removal of repressor, secondary structure in the 5’UTR regulates the initiation rate by changing the tim needed to complete an initiation event. Final protein output depends on both burst frequency and amplitude.

### Secondary structures in the 5’UTR and ribosome scanning

In the scanning model of translation initiation, the eIF4F complex binds the 5’-m^7^G cap and recruits the 43S pre-initiation complex to the mRNA. The 43S must traverse any structure in the 5’UTR to find the appropriate start codon, at which point the 60S or large subunit joins to form the elongation competent 80S ribosome and begin protein production. Structure in the 5’UTR is a major determinant of protein output and initiation efficiency (Kozak, 1991). Stem-loops near the 5’-cap inhibit binding of initiation factors and 43S (Kozak, 1986, 1989). However, secondary structures distal to the cap also significantly affect translational output in reporter assays and genome-wide studies (Babendure et al., 2006; Iwasaki et al., 2016; Rubio et al., 2014; Wolfe et al., 2014). The mechanism for this inhibition remains unclear. Using defined secondary structures within the 5’UTR, our experiments sought to systematically investigate the role that altered scanning plays at the single mRNA level. When inserted RNA hairpins ∼150 nt downstream of the putative transcription start site and immediately prior to the Kozak sequence, the hairpins presented an increased barrier for ribosome scanning.

There are two models for how the small subunit associates with the cap-binding complex through scanning. In the cap-severed model, the cap is dissociated and as a result multiple scanning 43S ribosomes can build up in the 5’UTR. In the cap-tethered model, the cap and ribosome bound initiation factors travel with the scanning 43S through initiation, such that only one ribosome occupies the 5’UTR at a time. Using selective 43S ribosome profiling, cap-severed scanning has been observed in yeast (Archer et al., 2016) whereas cap-tethered scanning has been observed in mammals (Bohlen et al., 2020), but the mechanism has yet to be resolved in scanning 43S structures (Brito Querido et al., 2020).

If the cap-severed model holds true, we hypothesized that the hairpin secondary structure would control burst frequency (Figure 4A, Model A). When the secondary structure is folded, initiation is limited and 43S complexes form queues upstream of the hairpin; unfolding of the 5’UTR would then lead to a burst of translation. A prediction of this hypothesis is that when the hairpin is stronger, the inactive period would be longer, thus reducing protein production. However, the experimental data are not consistent with this model of temporal control. Instead, hairpins control the rate of ribosome loading and therefore the amplitude of translation bursts (Figure 4A, Model B). Protein output is largely explained by the number of ribosomes loaded during the active state (Figure 4). This model is consistent with a cap-tethered mechanism where each individual ribosome must traverse the 5’UTR alone for successful initiation each time. The stronger the hairpin, the longer it takes for 43S to unwind structure, and the less initiation occurs per unit time.

Recent single molecule experiments suggest scanning is fast relative to large subunit joining, and secondary structure near the start codon reverses the 43S scanning complex, sending it back in the 5’-direction prior to GTP-hydrolysis by eIF2 (Wang et al., 2021a). In this model, the repeated kinetic competition between hairpin re-folding and 60S joining is the basis for different initiation efficiencies, as stronger hairpins increase the frequency of 43S reverse displacement prior to the completion of initiation. This agrees with our observation that the secondary structure reduced the initiation rate on active mRNA. In fact, 43S footprints in the 5’UTR are depleted in the region immediately upstream of the start codon where hairpins are most likely to be found in human 5’UTRs (Bohlen et al., 2020; Wang et al., 2021a) and where our hairpins were placed, suggesting amplitude control may represent a widespread form of initiation regulation.

### mTOR signaling and bursting

mTOR is the major regulator of global translation response to a variety of stimuli including nutrient and energy availability. mTOR-signaling primarily exerts control over translation initiation through its regulation of 4E-BPs (Ma and Blenis, 2009). Importantly, the response to changes in mTOR signaling are not uniform across all mRNA species. TOP mRNAs are acutely sensitive to changes along this signaling axis (Hsieh et al., 2012; Meyuhas and Kahan, 2015; Thoreen et al., 2012). In the steady state, a reporter TOP mRNA (EEF1A) exhibited more “bursty” behavior evidenced by shorter burst and longer dwell times, when compared with a reporter lacking the TOP motif (Figure 5). Upon mTOR inhibition with Torin-1, the intrinsic dwell time increased and initiation rate decreased significantly, while the intrinsic burst time remained largely unchanged. Importantly, the TOP mRNA was more acutely affected for each of these parameters (Figure 6). These results demonstrated that cap-adjacent sequences can mediate the bursting frequency of mRNAs and that mTOR modulates both bursting frequency and amplitude.

### Nature of the active and inactive states

In our model, translation initiation cannot occur in the inactive state. In the active state, translation is allowed, but does not necessarily occur (for example, when the initiation is slow). We describe a simple two-state model defined by three parameters (*τ*_*B*_, *τ*_*D*_ and *k*_*i*_) that correspond to the intrinsic state switching parameters and initiation rate.

What is the nature of the active and inactive states? One possibility is that active eIF4F assembled on the cap represents the active state. There, the cap-bound eIF4F is tethered to the scanning 43S and does not dissociate from the cap after initiation and multiple rounds of initiation can occur during the active state. If eIF4F falls off or is disassembled, the mRNA becomes inactive. Another possibility is that the inactive state is determined by the binding of *trans*-acting repressive factors to the cap or 5’UTR. For example, the alternative cap-binding protein 4EHP (eIF4E2) implicated in ribosome associated quality control (Hickey et al., 2020) and miRNA-mediated silencing (Chapat et al., 2017) inhibits canonical initiation. For a TOP mRNA, the cap-binding protein LARP1 has been implicated as a key *trans*-acting factor (Fonseca et al., 2018). Structural studies showed that LARP1 binds preferentially to 5’-cap structures with a TOP motif, preventing the assembly of eIF4F (Lahr et al., 2017; Philippe et al., 2018). Transcriptome wide ribosome profiling identified a suite of LARP1-repressed transcripts with 5’TOP motifs, including EEF1A, sensitive to mTOR inhibition by Torin-1 (Philippe et al., 2020). Although a two-state model explained many of our measurements, we do not rule out the possibility that there exist multiple active or inactive states, just as for transcription (Rodriguez et al., 2019). It is possible that both positive and negative *trans*-acting factors may modulate these states. Future development of single molecule translation imaging, including the imaging of endogenous transcripts, may be required to identify these multiple translational states and to parse the pleiotropic effects of these *trans*-acting factors.

## Limitations

The SINAPs system relies on a series of tandem epitopes and RNA stem-loops to visualize both the TLS and mRNA. We tethered mRNAs to plasma membrane for long term tracking using TIRF illumination. There are limitations for this method. First, the evanescent field decreases exponentially in the axial direction. The positions of nascent peptides fluctuate in axial due to the nature of the flexible polymer, which causes heterogeneous brightness in TIRF field. As such, there may be experimental heterogeneity in the TLS intensity for live cell experiments. For this reason, during the live imaging experiments, we chose to only classify whether the translation is active or not, without detailed analysis of intensity value. Instead, we used fixed cell smFISH-IF and 3D imaging to calculate the integrated intensity of TLS and single proteins, which misses temporal information. Second, mis-localizing mRNA to the membrane from its native context may alter aspects of its metabolism including both translation and decay though we do not observe differences at the ensemble level. Third, the exogenous reporters used in this study may not undergo the same type of physiological processing as for an endogenous transcript.

## Acknowledgements

This study is supported by National Science Foundation (MCB 1817447), National Institutes of Health (R01GM136897) and Pew Charitable Trust (00030601) to B.W. and National Science Foundation (2105837 and 2148534) to J.L.. N.M.L. is supported by NIH training grant T32 GM007445. J.A.S. is supported by a NRSA F30 Fellowship from the National Cancer Institute (F30CA260910). N.K.S. is supported by the Jane Coffin Childs Memorial Fund for Medical Research. Dr. Luke Lavis at the Janelia Research Campus for the fluorescent dyes used in this study. Dr. Andrew Holland and Dr. Robert Singer for providing our parent U2-OS cell lines. Dr. Sergi Regot for providing Torin-1. We thank Dr. Daniel Goldman and Dr. Malgorzata Latallo for critical discussions and feedback.

## Author Contributions

Conceptualization, N.M.L., J.K., B.W.; Methodology, N.M.L, J.K., O.V., J.L., B.W.; Software, N.M.L, J.K., O.V., B.W.; Validation, N.M.L., J.K., J.A.S., N.S.; Formal Analysis N.M.L, J.K., O.V., J.A.S., N.S.; Investigation, N.M.L., J.K., J.A.S., N.S., O.V., P.R., B.N., C.W.; Resources; B.W., T.H., J.L., R.G.; Data Curation, N.M.L., J.K.; Writing – Original Draft, N.M.L., J.K.; Writing – Reviewing & Editing, All Authors; Visualization, N.M.L., J.K.; Supervision, B.W., J.L., T.H. R.G.; Project Administration, B.W., N.L., J.K.; Funding Acquisition B.W., T.H., J.L., R.G.

## Supplemental Figures and Movies

**Figure S1.**
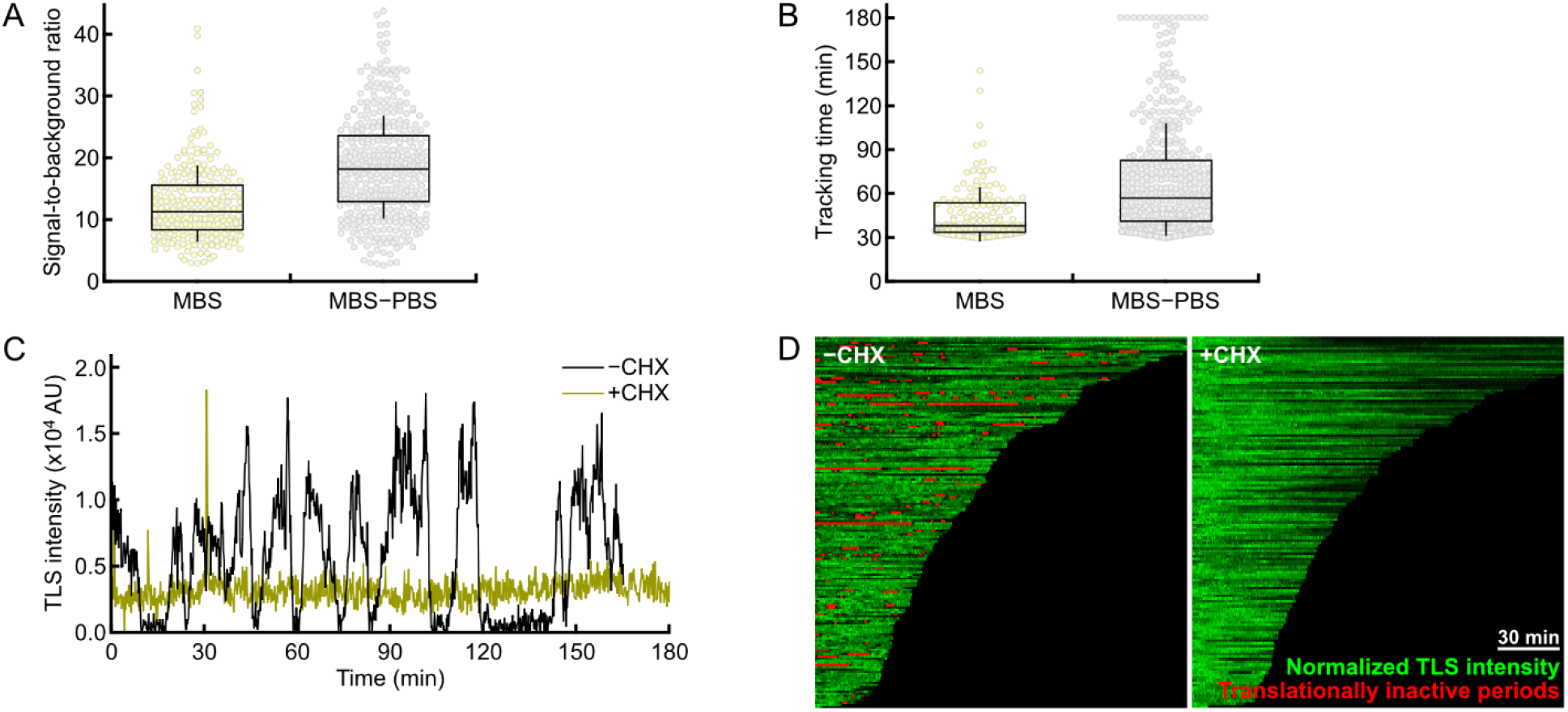
Membrane tethering comparison for translation site observation, associated with Figure 1. (A) Signal to background comparison between MBS and MBS-PBS tethering systems (n=150-439 mRNAs, box plots represent median ± interquartile range, whiskers for SD). (B) Raw tracking time comparison data for MBS and MBS-PBS tethering systems (n=150-439 mRNAs, box plots represent median ± interquartile range, whiskers for SD). (C-D) 100 µM cycloheximide (CHX) treatment 5 minutes prior to imaging ablates bursting activity. (C) Sample intensity traces with and without cycloheximide treatment. (D) Composite of single mRNA tracks for ST-AID reporter ± CHX (n=141-168 mRNA). MBS: MS2 binding site; PBS: PP7 binding site; SD: standard deviation.

**Figure S2.**
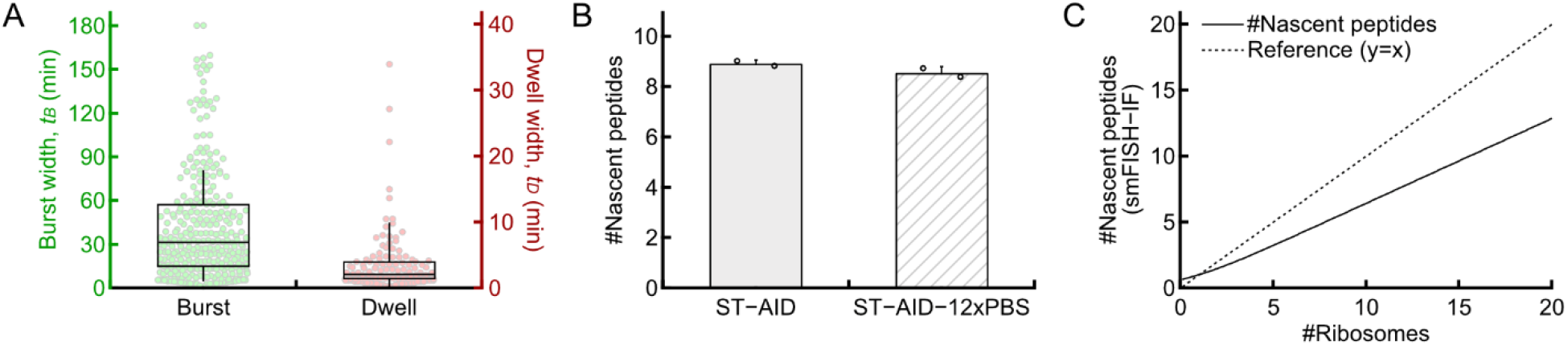
Burst measurement detail, associated with Figure 2. (A) Raw burst and dwell measurements for base ST-AID reporter (n=168 mRNA, box plots represent median ± interquartile range, whiskers for SD). (B) ST-AID-24xMBS and ST-AID-24xMBS-12xPBS show no difference in ribosome density as measured by smFISH-IF (n=627-1,037 TLS, mean of 2 biological replicates ± SD, ***p*** > 0.05 from Student’s t test). (C) Relationship between the measured number of nascent peptides from smFISH-IF and potential actual number of ribosomes. The NAP values underestimate the number of ribosomes in the experimentally measured range due to partially synthesized SunTag epitopes still translating the SunTag region (see STAR Methods). MBS: MS2 binding site; PBS: PP7 binding site; SD: standard deviations.

**Figure S3.**
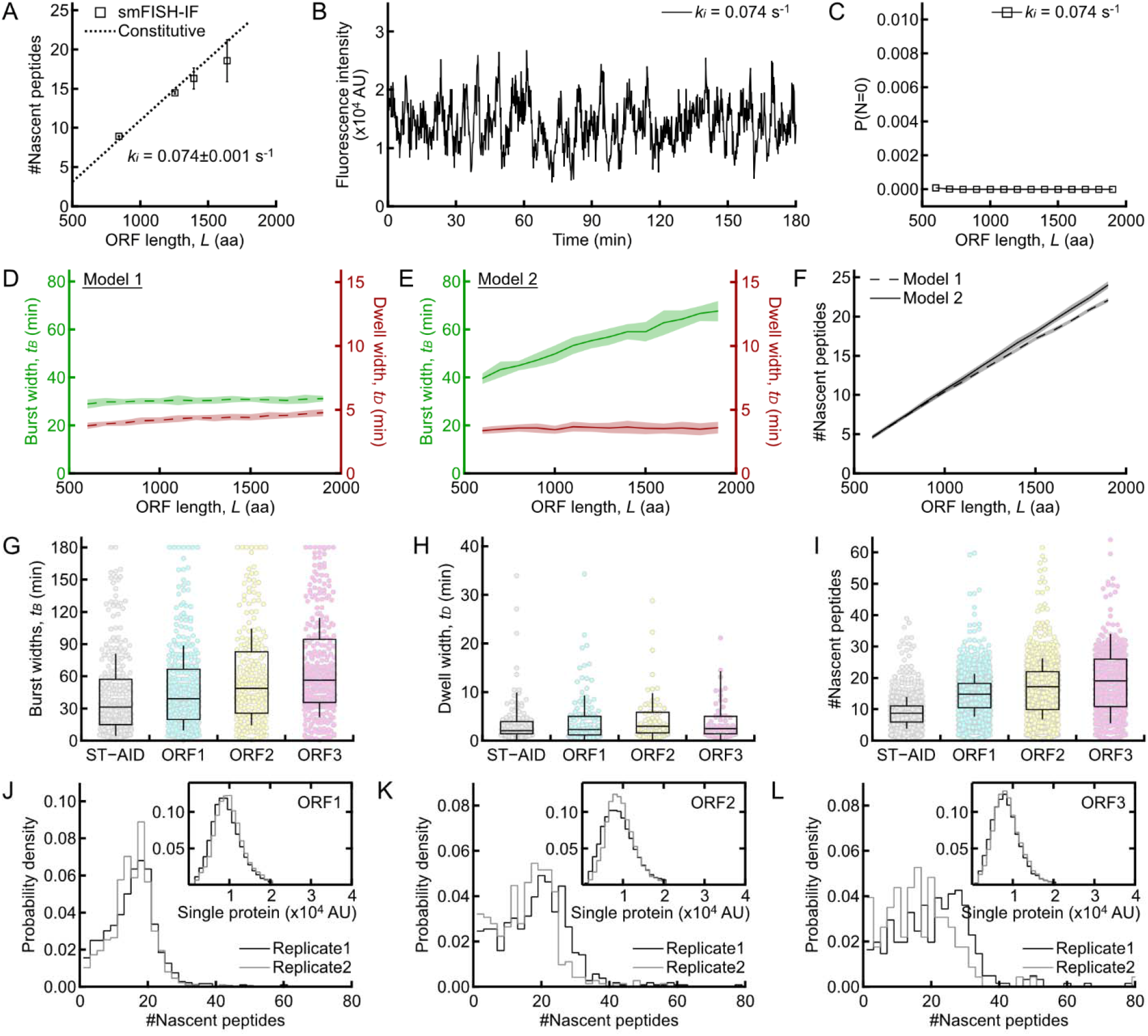
A two-state model for translational bursting, associated with Figure 3. (A) Nascent peptide density under constitutive initiation model. Experimental nascent peptide data from Figure 3E. (B) Sample simulated fluorescence intensity trace under constitutive initiation model. (C) Probability of observing zero ribosomes due to low initiation rate and dependence on ORF length is near zero at the initiation rate estimated from our observation. (D-F) Comparison of switching models through simulation. Model 1 (requiring ribosome run-off) does not show strong ORF length dependence of burst times. Model 2 (stochastic switching) demonstrates strong ORF length burst measurement dependency while leaving the dwell period unchanged. Simulation data plotted as mean ± SD. (F) Model 1 and 2 yield similar results for ribosome number and initiation rate. There is little contribution from bursting on overall ribosome density. Simulation data plotted as mean ± SD. (G-I) Raw burst (n=237-350), dwell (n=65-124) and NAPs density (n=419-1,291 TLS) data for ORF series reporters. Box plots represent median ± interquartile range (whiskers for SD). (J-K) Replicate single protein (insert) and normalized NAPs for ORF1-3 reporters (n=419-1,291 TLS, 29,065-43,939 single proteins). ORF: open reading frame; NAPs: nascent peptides; TLS: translation site; SD: standard deviation.

**Figure S4.**
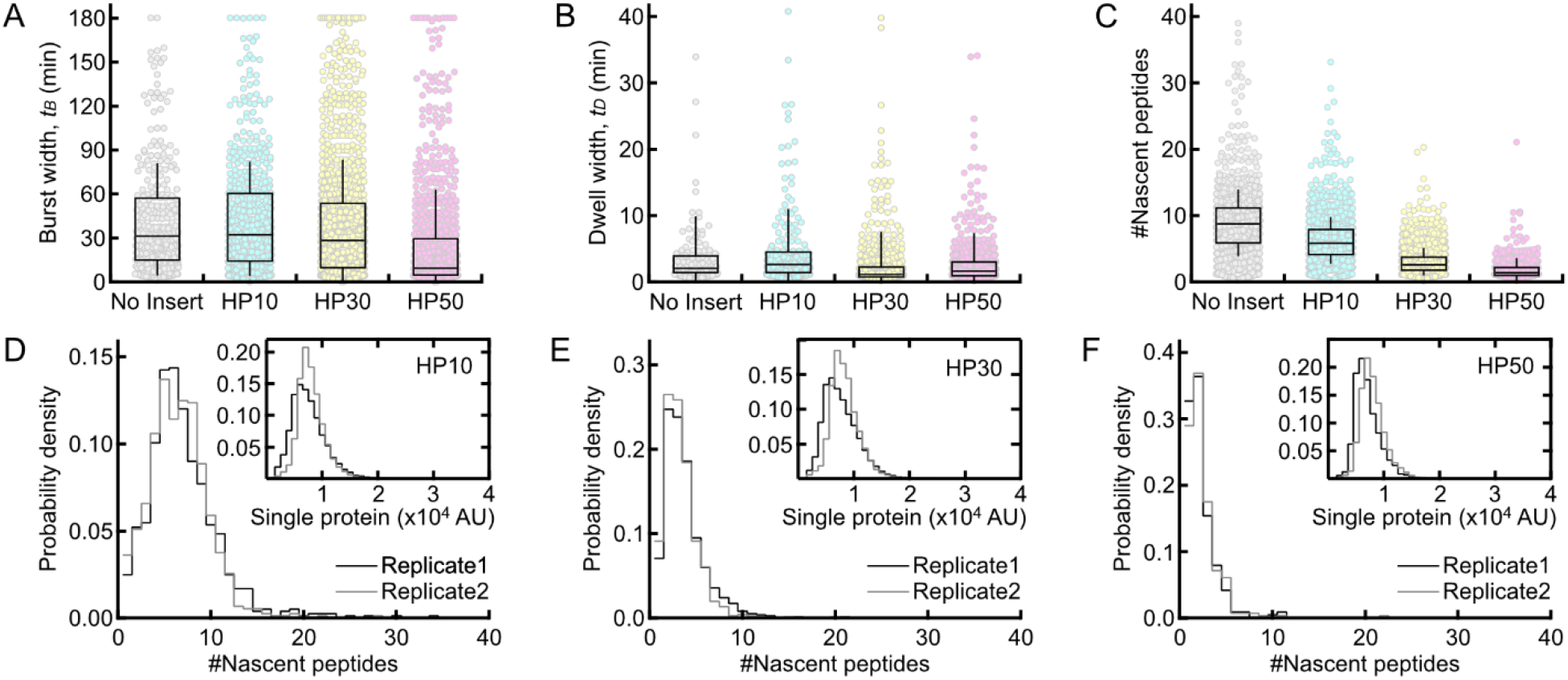
Hairpins alter burst amplitude, associated with Figure 4. (A-C) Raw burst (n=292-1,114), dwell (n=124-531) and nascent peptide measurements (n=493-1,509 TLS) for hairpin series reporters (box plots represent median ± interquartile range, whiskers for SD). (D-E) Replicate information for hairpin series for single protein intensity (insert) and normalized NAPs (n=493-1,509 TLS, 7,293-45,422 single proteins). NAPs: nascent peptides; TLS: translation site; SD: standard deviation.

**Figure S5.**
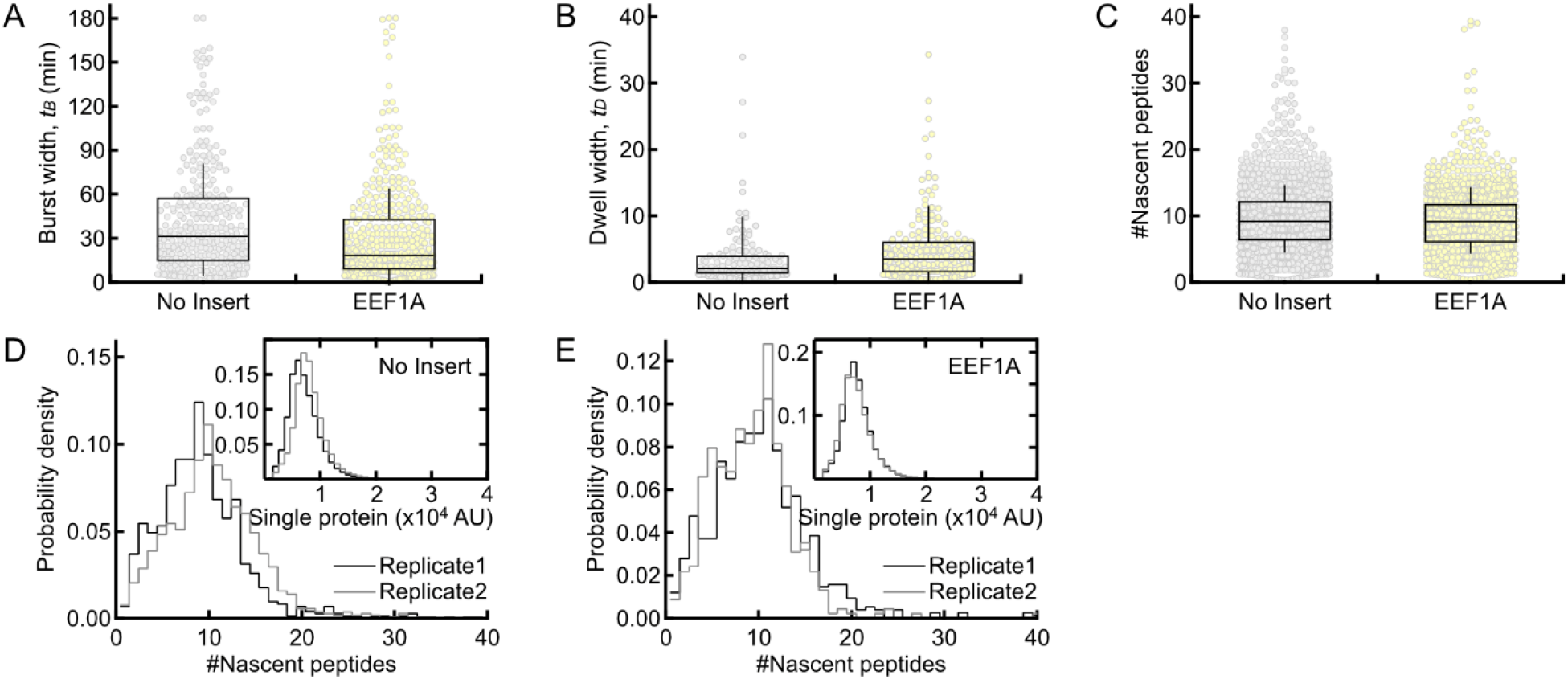
EEF1A TOP sequence confers unique burst characteristics, associated with Figure 5. (A-C) Raw burst (n=292-410), dwell (n=124-201), and NAPs measurements (n=1,037-1,205 TLS) for No Insert and EEF1A reporters (box plots represent median ± interquartile range, whiskers for SD). (D-E) Replicate information for No Insert and EEF1A single protein (inset) and NAPs (n=493-1,509 TLS, 41,228-56,672 single proteins). NAPs: nascent peptides; TLS: translation site; SD: standard deviation.

**Figure S6.**
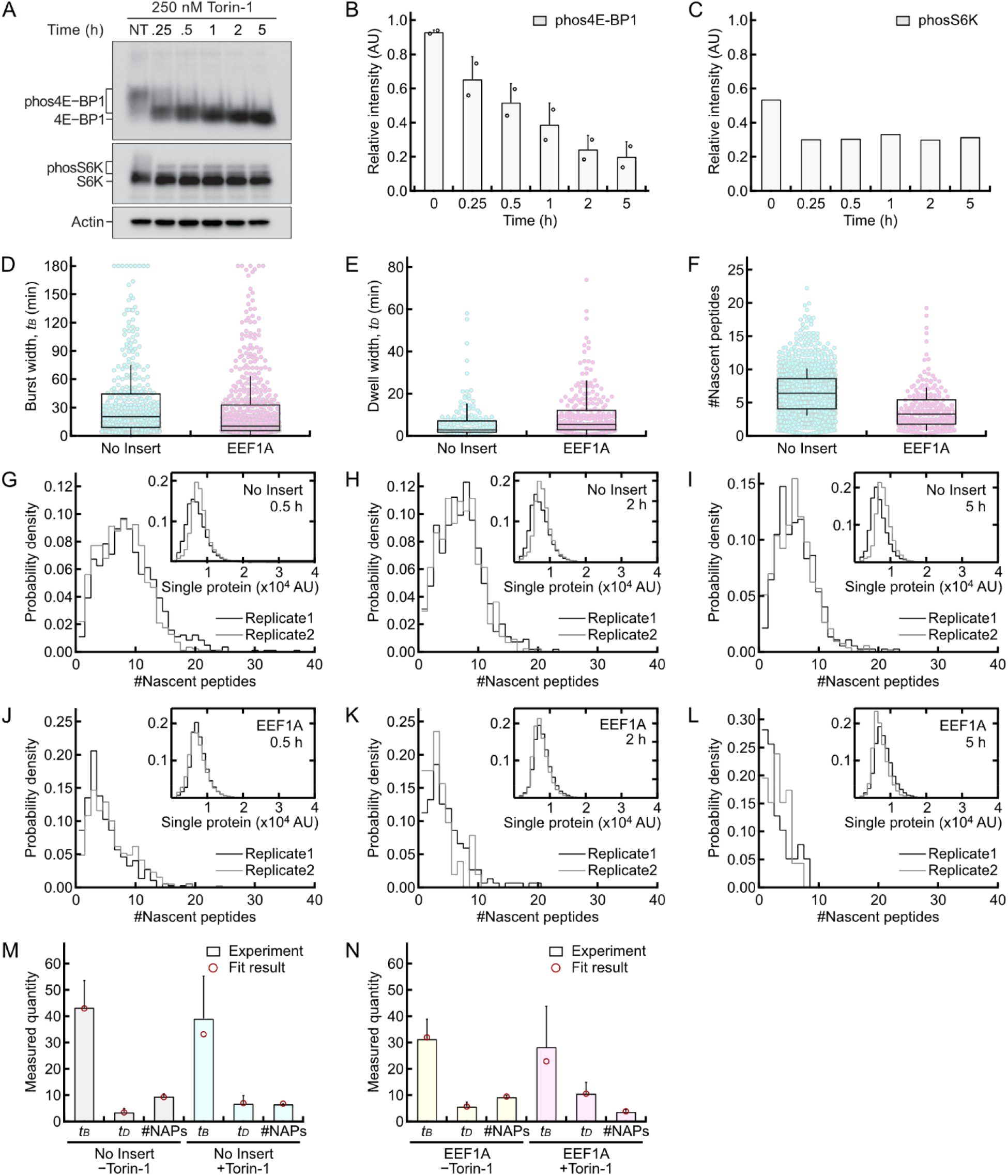
Torin treatment alters bursts characteristics for both No Insert and EEF1A reporters, associated with Figure 6. (A-C) Phos-Tag gel quantification of 250 nM Torin-1 response in U-2 OS imaging cell line (data represent 1-2 biological replicates ± SD). Phosphorylated 4E-BP levels stabilize between 2-5 hours (B) whereas S6K response reaches full response within 15 minutes (C). (D-F) Raw burst (n=377-545), dwell (n=193-249) and NAPs measurements (n=202-984 TLS) for No Insert and EEF1A reporters following 2 hours of 250 nM Torin-1 treatment (box plots represent median ± interquartile range, whiskers for SD). (G-L) Replicate information for No Insert and EEF1A single protein (insert) and NAPs following 250 nM Torin-1 treatment (n=202-984 TLS, 21,817-64,782 single proteins). (M-N) Phase diagram fit results for No Insert and EEF1A before and after Torin-1 treatment (single cell mean ± SD). NAPs: nascent peptides; TLS: translation site; SD: standard deviation.

**Table S1.** Summary on the measured burst frequency and amplitudes together with the extracted intrinsic properties from phase diagram analysis for all constructs used in this study, related to Figure 3-6 and S2-6. ^a^Values are presented as mean ± SD. ^b^Values are presented as fit result (lower bound, upper bound) obtained from the global fits using phase diagram. ***t***_***B***_: measured burst widths; ***t***_***D***_: measured dwell widths; #NAPs: number of nascent peptides; ***τ***_***B***_: intrinsic burst widths; ***τ***_***D***_: intrinsic burst widths; ***k***_***i***_: initiation rate; SD: standard deviations.

| Construct | #Cells | #mRNAs | $t_B$ (min)<sup>a</sup> | $t_D$ (min)<sup>a</sup> | #NAPs<sup>a</sup> | $\tau_B$ (min)<sup>b</sup> | $\tau_D$ (min)<sup>b</sup> | $k_i$ (min<sup>-1</sup>)<sup>b</sup> |
| --- | --- | --- | --- | --- | --- | --- | --- | --- |
| ST-AID | 8 | 168 | 43.3 ± 10.3 | 3.5 ± 1.5 | 8.9 ± 0.1 | 42 (36, 60) | 3.3 (2.8, 3.8) | 4.3 (4.3, 4.5) |
| ORF1 | 11 | 227 | 50.6 ± 9.0 | 3.8 ± 1.6 | 14.5 ± 0.2 |  |  |  |
| ORF2 | 7 | 172 | 57.7 ± 10.8 | 4.1 ± 2.5 | 16.3 ± 1.9 |  |  |  |
| ORF3 | 7 | 235 | 67.0 ± 13.6 | 5.1 ± 1.9 | 18.6 ± 3.8 |  |  |  |
| ST-AID | 8 | 168 | 43.3 ± 10.3 | 3.5 ± 1.5 | 8.9 ± 0.1 | 48 (36, 70) | 3.3 (2.8, 4.3) | 4.5 (4.5, 4.5) |
| HP10 | 13 | 315 | 47.1 ± 15.0 | 4.1 ± 1.6 | 6.3 ± 0.3 |  |  | 3.3 (3.3, 3.3) |
| HP30 | 17 | 643 | 43.2 ± 9.4 | 2.7 ± 1.5 | 3.1 ± 0.3 |  |  | 1.8 (1.8, 1.8) |
| HP50 | 9 | 300 | 29.3 ± 12.9 | 3.0 ± 1.2 | 1.9 ± 0.1 |  |  | 0.8 (0.8, 0.8) |
| ST-AID -Torin-1 | 8 | 168 | 43.3 ± 10.3 | 3.5 ± 1.5 | 9.5 ± 1.1 | 48 (24, 67) | 3.0 (3.0, 3.5) | 4.5 (4.5, 4.5) |
| ST-AID +Torin-1 | 8 | 184 | 39.1 ± 16.2 | 6.8 ± 3.1 | 6.6 ± 0.2 |  | 6.5 (5.5, 7.5) | 3.5 (3.3, 3.5) |
| EEF1A -Torin-1 | 6 | 209 | 31.4 ± 7.5 | 5.7 ± 1.7 | 9.3 ± 0.6 | 30 (22, 48) | 5.5 (5.0, 6.0) | 5.0 (4.8, 5.0) |
| EEF1A +Torin-1 | 10 | 296 | 28.3 ± 15.9 | 10.6 ± 4.3 | 3.7 ± 0.9 |  | 13 (12, 15) | 2.0 (2.0, 2.3) |

**Table S2.** STv4 smFISH oligonucleotide probe sequences, related to STAR Methods.

**Video S1**. ST-AID long-term translation imaging.

**Video S2**. EEF1A-ST-AID long-term translation imaging.

**Video S3**. ST-AID long-term translation imaging following 2hr 250 nM Torin-1 treatment.

**Video S4**. EEF1A-ST-AID long-term translation imaging following 2hr 250 nM Torin-1 treatment.

## Methods

### Resource Availability Lead contact

All material requests should be directed to Bin Wu.

### Materials availability

Reagents and materials produced in this study are available from Bin Wu pending a completed Materials Transfer Agreement. Constructs will be made available on Addgene.

### Data and code availability

All custom code for single molecule tracking, smFISH-IF analysis, and theoretical modeling can be found at (https://github.com/binwulab/uLocalize.git). Described in detail in Quantification and Statistical Analysis.

### Experimental Model and Subject Details

#### Cell lines and culture conditions

U-2 OS and HEK293T cells were cultured in DMEM (Corning, 10-013-CV) supplemented with 10% FBS (Millipore Sigma, F4135-500ML) and 100 units penicillin and 0.1 mg/mL streptomycin (Millipore Sigma, P0781) and maintained at 37 °C and 5% CO_2_. Cells were passaged every 2-3 days once they reached ∼75% confluency. Cells were tested monthly for mycoplasma infection and have always been negative.

#### Generation of stable cell lines

SINAPs auxiliary proteins including MCP/PCP variants, OSTIR1, and GCN4-scFv-sfGFP were stably expressed in U-2 OS cells using lentiviral transduction. tdMCP-HaloTag-IRES-tdPCP-SNAPtag-CAAX, tdMCP-HaloTag, or OSTIR1-IRES-scFv-sfGFP constructs in the lentiviral vector were transfected into HEK293T cells along with viral packaging components. 48 hours following transfection, the supernatant was collected, centrifuged to removed cell contents, and filtered through a 0.45 µm PVDF filter (Millipore Sigma, SLHV013SL). The supernatant was then concentrated in centrifugal filter unit (Amicon, UFC9100) and flash frozen in liquid nitrogen. Aliquots were thawed and diluted in 900 µL OptiMEM media (Gibco, 31985062) supplemented with 6 ng/mL polybrene (EMD Millipore, TR-1003-G) prior to application to cells. Cells were first infected with OSTIR-IRES-GCN4-scFv-sfGFP, expanded, and sorted for high GFP expressing cells. Following sorting, cells were infected with RNA coat proteins and sorted for positive cells.

Two cell lines were generated. First, live cell imaging lines were generated using a U-2 OS cell line harboring a Flp-In site and stably expressing a ponasterone A inducible system (Agilent), a gift from Robert Singer, were infected with tdMCP-HaloTag-IRES-tdPCP-SNAPtag-CAAX and OSTIR1-IRES-scFv-sfGFP. Cells were first sorted for high expression of scFv-sfGFP and high expression of tdMCP-HaloTag-IRES-tdPCP-SNAPtag-CAAX (cells were labeled with JFX-549 prior to sorting, see “Live cell translation imaging”). For data generated in Figure 5, a clonal line derived from this sort was used. For fixed cell imaging, a U-2 OS cell line harboring a Flp-In site and stably expressing the T-Rex tet-On system (Thermo Fisher), a gift from Andrew Holland, were infected with tdMCP-HaloTag and OSTIR1-IRES-scFv-sfGFP. Cells were sorted for high expression of scFv-sfGFP and low expression of tdMCP-HaloTag (cells were labeled with JFX-549 prior to sorting, see “Live cell translation imaging”).

To generate cells stably expressing our mRNA reporters, 10 µg of each respective reporter in the pcDNA5 vector backbone was restriction digested overnight with FspI to linearize the plasmid. Following restriction digest, the reporters were ethanol precipitated and resuspended in 10 µL of water and the concentration was measured. 1 µg of digested plasmid was added to the bottom of a 1.5 mL tube and 800,000 cells suspended in 20 µL SE nucleofection solution (16.4 µL nucleofection solution, 3.6 µL nucleofection supplement, Lonza, V4XC-1012). The solution was then transferred to a well in a 16-well Nucleocuvette™ Strip and nucleofected in a Lonza 4D-Nucleofector™ running the CM-104 program for U-2 OS cells. Following nucleofection, 100 µL of culture media was added to each nucleofection well and cells were plated in a 6-well dish. 800,000 U-2 OS cells that did not undergo nucleofection were also plated as a negative control. 48 hours following nucleofection, cells were trypsinized and re-plated in media containing 100 µg/mL hygromycin (InVivogen, ant-hg-1) to begin selection. Individual colonies appeared 7-10 days following transfection. Cells were removed from hygromycin following complete cell death on the negative control plate.

### Method Details

#### Plasmids

The reporters from this study were derived from our original SINAPs reporter (Wu et al., 2016) with the modifications discussed in (Goldman et al., 2021) aimed at reducing cryptic splice isoforms. This base construct, pcDNA5_CMV_ST, contains a Nano Luciferase, BFP, and AID downstream of the SunTag in the ORF (1,254 codons). The 3’UTR contains 24xMBSv5. We cloned the pcDNA5_CMV_ST SINAPs reporter into a new pcDNA5 backbone lacking an FRT site and containing a start codon in advance of the hygromycin resistance gene to facilitate stable cell line generation. This new reporter series is called pcDNA5-tet2-ST. For live cell reporters containing 12xPBSv5, we inserted the PP7 binding sites (PBS) downstream of the 24xMBSv5 array.

To vary ORF length, we inserted variable length sequences between the SunTag array and the AID. These ORFs were derived from our original SINAPs studies (Wu et al., 2016). We generated 3 additional reporters pcDNA-tet2-ST-AID (841 codons), pcDNA-tet2-ST-FLuc-AID (1,393 codons), and pcDNA-tet2-ST-Fluc-BFP-AID (1,639 codons). In order to add hairpins in the 5’UTR, we inserted sequences corresponding to the -10 kcal/mol, -30 kcal/mol, and -50 kcal/mol hairpins developed in (Babendure et al., 2006) immediately upstream of the Kozak consensus sequence ahead of the start codon.

To create pHR-NLS-tdMCP-HaloTag-IRES-tdPCP-SNAPtag-CAAX, we first eliminated the HA-tag in phage-UBC-NLS-HA-tdMCP-HaloTag (Addgene, 104098) and inserted IRES-tdPCP-SNAPtag-CAAX downstream of the tdMCP-HaloTag. We then replaced the transgene content of pHR-scFv-GCN4-sfGFP-GB1-dWPRE (Addgene, 60907) with NLS-tdMCP-HaloTag-IRES-tdPCP-SNAPtag-CAAX to generate pHR-NLS-tdMCP-HaloTag-IRES-tdPCP-SNAPtag-CAAX which we used to generate lentiviral particles for the RNA coat proteins. We used the pubc-OSTIR1-IRES-scFv-sfGFP-nls plasmid from our original publication (Wu et al., 2016).

#### Microscope

Live cell data were acquired on a custom inverted wide-field Nikon Eclipse Ti-E microscope equipped with three Andor iXon Ultra DU897 EMCCD cameras (512×512 pixels), apochromatic TIRF 100x oil immersion objective lens (1.49 NA, Nikon, MRD01991), linear encoded XY-stage with 150 micron travel range piezo z (Applied Scientific Instrumentation), and LU-n4 four laser unit with solid state 405 nm, 488 nm, 561 nm, and 640 nm lasers (Nikon), a TRF89901-EM ET-405/488/561/640nm laser quad band filter set for TIRF applications (Chroma), and Nikon H-TIRF system. The x-y pixel size was 160 nm.

Fixed cell data were acquired on a custom wide-field inverted Nikon Ti-2 wide-field microscope equipped with 40x oil immersion objective lens (1.3 NA, Nikon, MRH01401), Spectra X LED light engine (Lumencor), and Orca 4.0 v2 sCMOS camera (Hamamatsu). 1.5x internal magnification was used to bring the total magnification to 60x. The x-y pixel size was 108.3 nm and the z-pixel size was 300 nm. Nikon Elements was used to control both microscopes.

#### Live cell translation imaging

150,000 U-2 OS cells stably expressing tdMCP-Halo-IRES-tdPCP-SNAPtag-CAAX and OSTIR-IRES-scFV-sfGFP were seeded in 35 mm glass-bottom dishes (Cellvis, D35-20-1.5-N) approximately 48 hours prior to imaging. The following day, media was exchanged for 2 mL of culture media supplemented with 500 µM 3-indoleacetic acid (IAA, Millipore Sigma, I2886). Each dish was transfected with 400 ng of the reporter plasmid using X-tremeGENE HP DNA transfection reagent (Roche, 6366236001), at a ratio of 400 ng DNA to 40 µL 37 °C serum-free DMEM to 1.6 µL reagent per dish being transfected. Reagents were added in that order and incubated at room temperature for 15 minutes and the solution was then added dropwise to the sample. 12-16 hours following transfection, cells were dyed by removing 100 µL of media, adding 4 µL of fresh IAA, and adding 2 µL of 10 µM JFX-549 (a gift from Luke Lavis) for a final concentration of 10 nM (Grimm et al., 2016, 2017). 30 minutes after adding dye, dishes were washed three times with warm culture media and then returned to incubator in culture media supplemented with 500 µM IAA. At the time of imaging, media was exchanged for Leibovitz-15 (L-15) media without phenol red (Gibco, 21083027) supplemented with 500 µM IAA and moved to the microscope.

Once on the microscope, the sample was warmed and maintained at 37 °C for the duration of the imaging. At the beginning of each imaging session, the laser was aligned to find a uniform 0 ° incidence angle before moving the excitation beam angle to 66 ° for TIRF imaging. Positively transfected cells containing 10-75 membrane tethered mRNAs and translation sites were identified for imaging as the samples came up to temperature equilibrium. 1-4 positions were identified for each imaging period and acquired in the same sequence. Cells were excited simultaneously with 488 nm (4% power) and 561 nm (5% power) lasers. The exposure time was 500 ms and data was acquired at a 10-s frame rate for 3 hours (1,081 frames total). For samples that received cycloheximide treatment, positions were identified and 1 mL of imaging media was removed from the dish. Cycloheximide was added to a final concentration of 100 µg/mL (in the total imaging volume) and added dropwise back to the sample on the microscope and mixed. 5 minutes following treatment, imaging began. For samples undergoing treatment with Torin-1 (SelleckChem, S2827), media was exchanged for L-15 media without phenol red supplemented with both 500 µM IAA and 250 nM Torin-1. Positions were identified as the sample came up to temperature, but imaging did not commence until 2 hours following treatment.

#### smFISH probe labeling

smFISH probes were synthesized as outlined described in (Gaspar et al., 2017) with some modifications. 48 20-mer oligonucleotides (Table S2) were ordered in an array format from IDT, pooled, and conjugated to amino-11-ddUTP (Lumiprobe, A5040) at the 3’-end using terminal deoxynucleotidyl transferase (TdT) (Thermo Fisher, EP0162). Following size-exclusion purification in a Spin-X centrifuge column (Corning, 8161) loaded with Bio Gel P-4 Beads (Bio Rad, 1504124), the oligonucleotide-amino-11-ddUTP was labeled with Cy5-NHS ester (Lumiprobe, 43020). Following labeling, the probes were purified again through a Spin-X centrifuge column to remove non-conjugated dyes. Labeling efficiency and yield were determined by nanodrop.

#### smFISH coupled with immunofluorescence (smFISH-IF)

We performed smFISH-IF on U-2 OS cells stably expressing the mRNA reporters with 24xMBSv5 3’UTRs in cells. German glass 18 mm #1 coverslips (Electron Microscopy Services, 72292-09) placed in a 12-well dish were washed in 3 M sodium hydroxide (Millipore Sigma, 221465) for 30 minutes. Following etching, coverslips were coated with 2.5 µg/mL bovine plasma fibronectin (Millipore Sigma, F1141-2MG) diluted in phosphate buffered saline (PBS, Corning, 46-013-CM). The coverslips were washed 4x with PBS, and 50,000 cells were plated per well in culture media. 24 hours following plating media was exchanged and supplemented with 250 µM IAA.

1 hour prior to fixation, the culture media was supplemented with 2 µg/mL doxycycline hyclate (Millipore Sigma, D9891) with fresh 250 µM IAA. smFISH-IF was done as described in (Latallo et al., 2019). All solutions were prepared in nuclease free water (Quality Biological, 351-029-131CS). Cells were washed 3x with 1xPBS with 5 mM magnesium chloride (PBSM, Millipore Sigma, M2670-500G). Cells were then fixed for 10 minutes at room temperature in PBSM supplemented with 4% paraformaldehyde (Electron Microscopy Sciences, 50-980-492). Cells were then washed 3x for 5 minutes in PBS and exchanged into a permeabilization buffer consisting of PBSM + 5 mg/mL BSA (VWR, 0332-25G) + 0.1% Triton-X100 (Millipore Sigma, T8787-100mL) and maintained for 10 minutes at room temperature. Following permeabilization, the samples were washed 3x for 5 minutes in PBSM and exchanged into a pre-hybridization buffer consisting of 2xSSC (saline-sodium citrate buffer, Corning, 46-020-CM), 10% formamide (Millipore Sigma, F9037-100ML), and 5 mg/mL BSA. The sample was then exchanged into a hybridization solution of 2xSSC, 10% formamide, 1 mg/mL competitor *E. coli* tRNA (Millipore Sigma, 10109541001), 10% w/v dextran sulfate (Millipore Sigma, D8906-100G), 2 mM ribonucleoside vanadyl complex (NEB, S1402S), 100 units/mL SUPERaseIn (Thermo Fisher, AM2694), 60 nM SunTag_v4-Cy5 smFISH probes, and chicken anti-GFP (1:1,000 dilution, Aves Labs, GFP-1010). The sample was incubated for 3 hours at 37 °C.

Following hybridization, the coverslips were washed 4x with PBS containing 10% formamide. The samples were then incubated for 20 minutes twice with a goat anti-chicken IgY secondary antibody labeled with Alexa Fluor 488 (Thermo Fisher, A-11039). The samples were further washed 3x with 2xSSC. A final 5-minute 2xSSC wash was performed before the coverslips were mounted on a pre-cleaned frosted glass slide (Thermo Fisher, 12-552-3) with ProLong Diamond antifade reagent with DAPI (Invitrogen, P36962).

#### Phos-tag gels measuring phosphorylated 4E-BP and phosphorylated S6K levels

For Phos-tag immunoblotting, cells were lysed in RIPA buffer (25 mM Tris-HCl pH 7.6, 150 mM NaCl, 1% NP-40, 1% sodium deoxycholate, 0.1% SDS, Thermo Fischer, 89900) supplemented with 1x Halt protease and phosphatase inhibitor cocktail (EDTA-free, Thermo Fisher, 78445), benzonase (50 units/mL, Millipore Sigma, E1014), 100 µg/mL anisomycin, 10 mM sodium pyrophosphate, 10 mM beta glycerophosphate, and 1 mM TCEP. Lysates were collected by cell scrapers and clarified by brief centrifugation at 8000 *g* (7 min, 4 □). The concentration of clarified supernatants was measured using BCA assay (Thermo Fisher, 23228, 23224). Samples were resolved by 7% (for p70 S6K) or 8% (for 4E-BP1) SDS-PAGE containing 10.7 µM Phos-tag (Wako, AAL-107) and 21.3 µM MnCl_2_, and transferred to PVDF membranes as per manufacturer’s instructions. Membranes were blocked in 5% non-fat dry milk resuspended in Tris-Buffered Saline (TBST, 1 h, 25 □) with gentle rocking, followed by overnight incubation with primary antibodies for 4E-BP1 (CST, 9644) or S6K (CST, 9202) in 5% non-fat dry milk in TBST at 4 □, followed by 4x 10 min washes in TBST at 25 □, followed by incubation with HRP-conjugated secondary antibody (1:5,000 dilution) in 5% non-fat dry milk in TBST (1 h, 25 □), followed by 4x 10 min washes in TBST. All incubation steps were performed with gentle rocking. Western blots were visualized by HRP chemiluminescence using Super Signal West HRP substrate (Thermo Fisher) using a BioRad ChemiDoc imager.

#### Quantifying reporter mRNA Levels with RT-qPCR

U-2 OS cells stably expressing the tdMCP-HaloTag-IRES-tdPCP-SNAPtag-CAAX and scFv-sfGFP constructs were seeded in duplicate for each condition to a density of 200,000 cells per well in DMEM supplemented with 10% FBS and 1x pen/strep in a 6-well dish. The following day, each well was co-transfected with 1 µg of NanoLuciferase (NLuc) expressing reporter plasmid containing no insert, a 10 kcal/mol hairpin (pcDNA-tet2-HP10-ST-NLuc-BFP-AID-24xMS2v12xPP7v5), a 30 kcal/mol hairpin (pcDNA-tet2-HP30-ST-NLuc-BFP-AID-24xMS2v5/12xPP7v5), or a 50 kcal/mol hairpin (pcDNA-tet2-HP50-ST-NLuc-BFP-AID-24xMS2v5/12xPP7v5), structure in the 5’UTR and 70 ng of firefly luciferase (FLuc)-expressing normalizer plasmid (pcDNA5_FRT_TO_EGFP_AID_Luciferase, a gift from Andrew Holland) using X-tremeGENE 9 DNA transfection reagent (Roche, 06365787001) at a ratio of 5 µL reagent to 1 µg DNA to 100 µL Opti-MEM reduced serum medium (Gibco, 31985062). Approximately 24 h later, cells were washed in 1 mL PBS and lysed directly in 1 mL of TRIzol (Thermo Fisher, 15596018) according to the manufacturer’s protocol. RNA was harvested and precipitated using isopropanol according to the manufacturer instructions. The resultant RNA pellet was resuspended in 40 µL of 1x TURBO DNAse buffer with 2 µL of TURBO DNAse (Thermo Fisher, AM2239) and incubated at 37 □ for 2 h at 37 □. DNAse was removed using the TURBO DNAse-free inactivation reagent (Thermo Fisher, AM1907) according to the manufacturer’s instructions.

Reverse transcription quantitative PCR (RT-qPCR) of the reporter constructs was performed as previously described (Goldman et al., 2021) with slight modifications. RNA was reverse transcribed using the ProtoScript II First Strand cDNA Synthesis Kit (NEB, E6560) using the random primer hexamer mix and 4 µL of RNA per 20 µL reaction. A no-RT control reaction was included for two samples substituting water for the enzyme as a control for amplification of genomic DNA. qPCR reactions were performed in a 384-well plate using iTaq Universal SYBR Green Supermix (BioRad, 1725121) in a 10 µL reaction with 1 µL template cDNA and 333 nM primer concentration. Primers were designed amplify only from un-spliced reporter mRNAs. Each qPCR reaction was performed in triplicate for each sample using primers for both NLuc (reporter plasmid) and FLuc (normalizing plasmid). A series of 8 2-fold dilutions was generated from one sample and probed with each primer pair to generate standard curves. qPCR was performed in a QuantStudio 6 Real-Time PCR System from Thermo Fisher.

#### Luciferase assay for quantification of reporter protein levels

U-2 OS cells stably expressing the tdMCP-HaloTag-IRES-tdPCP-SNAPtag-CAAX and scFv-sfGFP were seeded in sextuplicate in a 96-well plate in DMEM supplemented with 10% FBS and 1x pen/strep at a density of 5,000 cells per well. The following day, each well was transfected with 28 ng of NLuc-expressing reporter plasmid containing no insert, a HP10, a HP30, or a HP50 hairpin structure in the 5’UTR and 2 ng of FLuc-expressing normalizer plasmid (pcDNA5_FRT_TO_EGFP_AID_Luciferase, a gift from Andrew Holland) using X-tremeGENE 9 DNA transfection reagent (Roche, 06365787001) at a ratio of 5 µL reagent to 1 µg DNA to 100 µL Opti-MEM reduced serum medium (Gibco, 31985062). Approximately 24 h later, NLuc and FLuc activities measured using the NanoGlo Dual-Luciferase Reporter Assay System (Promega N1630) in a Synergy H1 microplate reader (BioTek). The ratio of NLuc to FLuc (in arbitrary units) was measured to determine relative protein output.

### Quantification and Statistical Analysis

#### Segmentation of live cell imaging data

Data were processed using a combination of custom MATLAB code, AirLocalize (Lionnet et al., 2011) and u-track (Jaqaman et al., 2008). Particle detection was performed using the AirLocalize gaussian mask algorithm (Lionnet et al., 2011; Thompson et al., 2002), while u-track was used for tracking in each channel. Additionally, we integrated a custom analysis module to link tracks in different channels within the u-track software to facilitate mRNA and translation site (TLS) colocalization. This custom module takes independently detected and tracked mRNA and TLS tracks and links them for downstream analysis. For periods of low translation or translational inactivity, the mRNA position is used as the location for the gaussian mask in the TLS channel, enabling the detection of weak TLS and measuring the background signal, thus interpolating the intensity trace between bursts. RNA tracks shorter than 30 minutes (180 frames) were discarded from co-localization analysis and co-localized translation tracks are required to last at least 50 seconds (5 frames) to be considered co-localized.

To facilitate data inspection, we generated the TrackViewer tool in MatLab. TrackViewer takes our colocalized mRNA and TLS tracks and displays tracking and movie data simultaneously for visual inspection. Tracks with inaccurate tracking or that overlap with other mRNAs or TLS are discarded from the data set, ensuring only accurate, co-localized tracks are analyzed. Following data inspection, we segmented tracks into translationally active and inactive periods. We first identify all extended periods in the track where the TLS signal drops below 5% of the maximum value of each track. Then using a combination of the fluorescence trace data and comparison with the movie in TrackViewer, we visualize and edit all transition points based on visual inspection.

Translational bursts are the length of all translation events that exceed 1 minute. Dwell periods all periods between the start of the mRNA track and translation or the time of inactivity between two translation events. Dwell periods are defined as inactive periods followed by the return of translation. Translationally inactive periods that do not return to the translating state are not considered in the dwell time distribution as they may represent events such as RNA degradation.

#### Modeling overlapping burst and dwell periods

With the two-state random telegraph model, the on- and off-transitions occur stochastically with the rates of *k*_*on*_ (on-switching rate) and *k*_*off*_ (off-switching rate), thus, the intrinsic burst and dwell widths (***τ***_***B***_ and ***τ***_***D***_) are exponentially distributed, and their probability density functions (*PDF*) can be described as below.

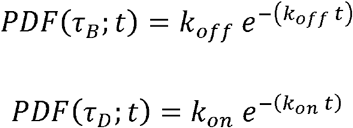

Experimentally measured burst and dwell widths (***τ***_***B***_ and ***τ***_***D***_) can be different to the intrinsic ones. When the mRNA is turned on, there might be a short time delay (*τi*) between the *τ*_*B*_ and *t*_*B*_ corresponding to the average initiation time (*τ*_*i*_ = 1/*k*_*i*_, *k*_*i*_ is the initiation rate). When the mRNA is turned off, it still can be observed for the average ribosome run-off time (*τ*_*RO*_ = *L*/*k*_*e*_, *L* and *k*_*e*_ are the ORF length and elongation rate, respectively) until there is no ribosome on it. Thus for the single *τ*_*B*_ and *τ*_*D*_, the mean widths of *t*_*B*_ and *t*_*D*_ can be described as below.

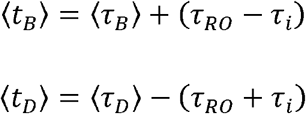

Here we assumed that *k*_*i*_ is sufficiently fast so that the fluorescent tag (SunTag-bound scFv-sfGFP in our measurement) persists during the intrinsic burst periods. Based on previously reported values of *k*_*i*_ and *k*_*e*_, *τ*_*i*_ is much shorter than *τ*_*B*_ and *τ*_*RO*_, we therefore assumed that *τ*_*i*_ can be negligible (Wu et al., 2016).

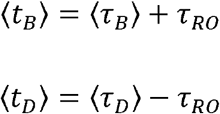

When an *τ*_*D*_ is shorter than *τ*_*RO*_, this dwell event cannot be observed and adjacent two bursts should be overlapped. The probability of -sequential overlap events, *P*(*OV*;*n*), is determined by the distribution of *τ*_*D*_ and *τ*_*RO*_.

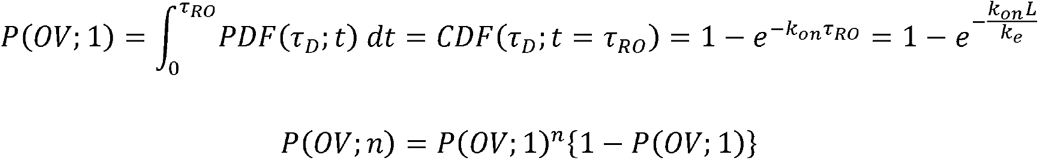

And the *n* sequential overlap events result in a measured burst, 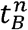, that consists of (*n* + 1) × *τ*_*B*_,

*n* ×*τ*_*D*_, and 1 ×*τ*_*RO*_. Since *τ*_*B*_ and *τ*_*D*_ follow the exponential distribution, (*n* + 1) × *τ*_*B*_ and *n* ×*τ*_*D*_ follow the gamma distributions with the means of (*n* + 1)/*k*_*off*_ and *n*/ *k*_*on*_, respectively. Thus, the mean width of a measured burst with overlap events can be described as below.

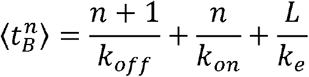

Taken together, the mean width of the *t*_*B*_ is described as below.

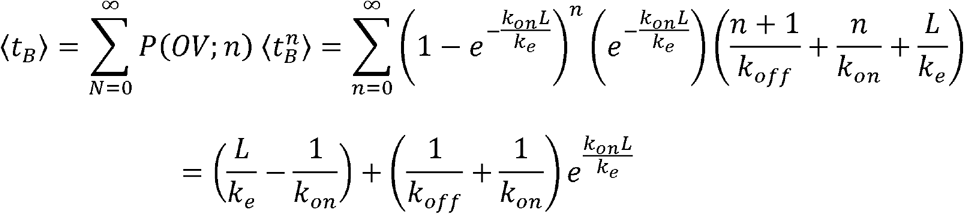

In the dwell width measurement, if an intrinsic dwell event is shorter than *τ*_*RO*_ then it is not detectable as the mRNA is turned on before the complete ribosome run-off. Therefore, *t*_*D*_ follows a truncated exponential distribution as below.

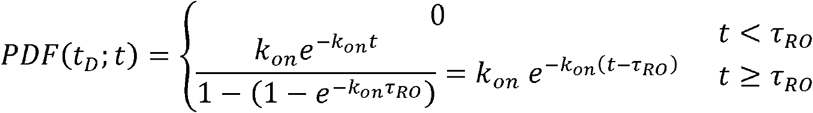

And *τ*_*D*_ longer than *τ*_*RO*_ should have reduced widths by *τ*_*RO*_.

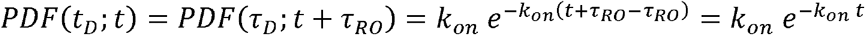

Thus, *t*_*D*_ should follow the same distribution to *τ*_*D*_, the exponential distribution with the same mean, but the frequency of observation is decreased by *CDF* (*τ*_*D*_;t = *τ*_*RO*_). To be commensurate with our experimental resolution and analysis (see above), we ignored *t*_*D*_ shorter than 3 frames (<30 s), which introduced another truncated distribution for the *t*_*D*_.

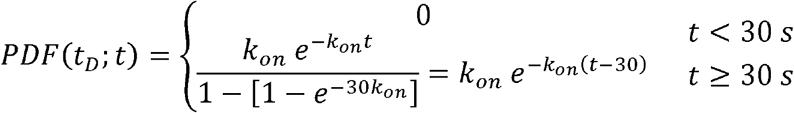

Mean value of *t*_*D*_, < *t*_*D*_>, can be calculated as below:

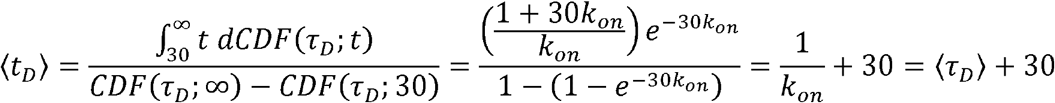

Thus, our measured dwell (*t*_*D*_) may show slightly increased value than the intrinsic dwells (*τ*_*D*_) due to the analysis process. Overall, by allowing overlap events between bursts, the measured burst width (*t*_*B*_) elongated with longer ORF lengths as the probability of overlap is increases, whereas the mean dwell width (*t*_*D*_) is not affected by the ORF length (Figure S3E).

#### Monte Carlo simulation of translation process

A homemade Python script was used to perform Monte Carlo simulation (Rossum and Drake, 2009). To simulate translation under the bursting, we utilized TASEP-based Monte Carlo method for the translation elongation process. With four input parameters (initiation rate *k*_*i*_, intrinsic burst width *τ*_*B*_, intrinsic dwell width *τ*_*D*_ and ORF length *L*), we obtained the three model outputs (burst width *t*_*B*_, dwell width *t*_*D*_ and number of nascent peptides #*NAPs*) by simulating fluorescence intensities on mRNA and applied the same analysis in experiment.

A brief workflow of the Monte Carlo simulation can be described as below.

i. We generated different track lengths over time (*L*_*T*_) for each mRNA to mimic our experimental condition. That is, the track lengths follow a truncated exponential distribution, whose lower limit, rate parameter, and upper limit are extracted from experimental measurements.
ii. Additional 100 frames (1 frame = 10 s) were simulated at the beginning of each mRNA to ensure all mRNAs are in the translational equilibrium.
iii. At the simulation time step (Δ*t*) of 0.02 s, the following simulations were performed to capture the stochastic nature of sub-translation processes. This small Δ*t* ensures that the probabilities of stochastic events (= Δ*t* × *k*_*event*_) are much less than 1 for given kinetic parameters (*k*_*on*_ << 1 s^-1^, *k*_*off*_ << 1 s^-1^, *k*_*i*_ << 1 s^-1^ and *k*_*e*_ = 4.7 codons/s).
  a. We determined the intrinsic status of mRNA by assuming the stochastic on- and off-switching events with the rates of *k*_*on*_ and *k*_*off*_, respectively.
  b. Ribosome initiation can occur stochastically only when the mRNA is intrinsically active.
  c. Existing ribosomes have a chance to move a single codon stochastically. When there are more than two ribosomes on a mRNA, the update order of ribosomes was determined randomly at each simulation step. We set a ribosome footprint as 9 codons per ribosome, so a ribosome cannot move further if there is another ribosome within 9 codons ahead, consistent with the measurement of ribosome footprints (Ingolia et al., 2009).
  d. If a ribosome is located at the end of the mRNA, it has a chance to leave the mRNA with a termination rate (*k*_*t*_= *k*_*e*_/2).
  e. For the remaining ribosomes, the number of fluorescent tags were calculated so that it is proportional to the sum of the travelled distance of each ribosome on the mRNA.
iv. The fluorescence intensities (*FI*) were generated from the number of fluorescent tags at each frame. The *FI* of single tag was extracted from the smFISH-IF measurement by assuming a single nascent peptide with 24-tags has a gamma-distributed *FI*.
v. Following analysis steps were applied to the simulated *FIs*.
  a. To re-estimate the #*NAPs* from the *FI*, we sampled the *FI* for every 60 frames to avoid short-term memory effect on *FI* measurement. Sampled *FI*s were divided by the mean value of single nascent peptide *FI*. We rejected the #*NAPs* below 0.2 (the same threshold with the experimental analysis).
  b. To binarize the *FI* traces, we first applied the median filter to smooth the traces. For each filtered *FI* traces, we set a threshold of 0.05×max(*FI*) to define the measured burst and dwell status of the target mRNA. If there were bursts or dwells shorter than 3 frames, we removed them and linked adjacent dwells (for burst removing) or bursts (for dwell removing) because those very short events most likely come from the noise of fluorescence intensity.

The phase diagram was generated for using the four input parameters as follows:

i. *k*_*i*_ (min^-1^): 0.50, 0.75, 1.00, 1.25, 1.50, 1.75, 2.00, 2.25, 2.50, 2.75, 3.00, 3.25, 3.50, 3.75, 4.00, 4.25, 4.50, 4.75, 5.00, 5.25, 5.50, 5.75, 6.00, 6.25, 6.50, 6.75, 7.00 (total 27 conditions).
ii. *τ*_*B*_ (min): 10.0, 11.9, 13.9, 15.6, 16.7, 17.9, 19.2, 20.0, 22.0, 24.0, 26.0, 28.0, 30.0, 32.0, 34.0, 36.0, 38.0, 40.0, 41.7, 44.0, 46.0, 48.0, 50.0, 51.9, 53.9, 55.9, 58.0, 60.0, 63.4, 66.7, 70.0 (total 31 conditions).
iii. *τ*_*D*_ (min): 1.00, 1.36, 1.67, 2.00, 2.25, 2.50, 2.75, 3.00, 3.25, 3.50, 3.75, 4.00, 4.25, 4.50, 4.75, 5.00, 5.50, 6.00, 6.50, 7.00, 7.50, 8.00, 8.50, 9.00, 9.50, 10.00, 10.50, 11.00, 11.50, 12.00, 12.50, 13.00, 13.50, 14.00, 14.50, 15.00 (total 36 conditions).
iv. *L* (codons): 841, 1254, 1393, 1639 (total 4 conditions).

The number of mRNAs simulated for each condition was 3,000 (*τ*_*B*_ ≤ 30) or 4,000 (*τ*_*B*_ > 30), and the means of *t*_*B*_, *t*_*D*_ and #*NAPs* were used to estimate the intrinsic properties by comparing with the experimental measurements.

#### Determining intrinsic burst properties with phase diagram analysis

To extract the intrinsic properties from the generated phase diagram, we fit the experimental results to the phase diagram to find a combination of input parameters that minimizes the root-mean-squared errors (*RMSE*) of measured values from experiment and simulation. Since we have three different measurements (*t*_*B*_, *t*_*B*_ and #*NAPs*), *RMSE* of each measurement was normalized by using the standard error of the means (*SEM*) from the experiment.

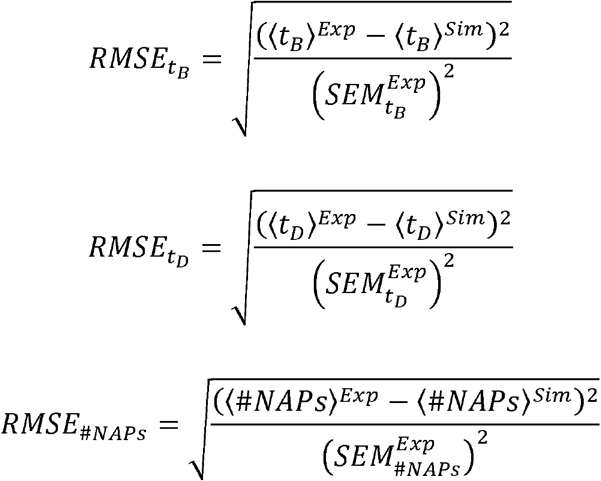

Superscribed *Exp* and *Sim* represents that the values come from the experiments and the simulation, respectively. The best combination of input parameters should minimize the differences of all three measurements, so we can get the intrinsic properties by minimizing *RMSE*_*All*_, the sum of *RMSE*s of each measurement.

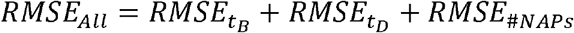

The confidence intervals (*CI*s) of this fitting were determined as the input parameters that give smaller differences between *RMSE*_*All*_ and the minimum value than a threshold, which is set to 1 for individual fit processes.

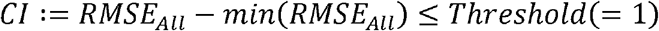

Individual fit for each construct showed heterogeneous intrinsic properties (data not shown), due to the different noise levels in different experimental measurements and/or that some of the experimental conditions may perturb the global state of the cell. To better interpret our observations, we performed a global analysis across the constructs within each experimental series by making some assumptions. For example, for the ORF-series (Figure 3 and Figure S3), we hypothesized that ORF length does not affect the intrinsic property to find a shared set of *τ*_*B*_, *τ*_*D*_ and *k*_*i*_ that can explain all the experimental measurements. Consequently, we found a global minimum of the sum of *RMSE*_*All*_ for the four ORF-series constructs, ST-AID, ORF1, ORF2, and ORF3.

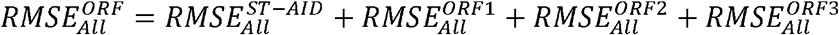

*CI*s of ORF-series were calculated in the same way, but we set the threshold as 4 instead of 1 because we added the *RMSE*s for all constructs.

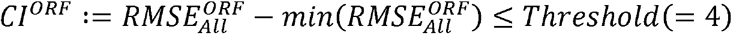

In the Model B of hairpin-series experiment (Figure 4 and Figure S4), we assumed that 5’UTR structure only affects the and all the constructs share the *τ*_*B*_ and *τ*_*D*_. Similarly, we assumed that Torin-1 treatment does not change *τ*_*B*_ and only alters *τ*_*D*_ and (Figure 6 and Figure S6). If a part of intrinsic parameters was assumed to be shared like in above examples, we first fit the non-shared parameters for all combinations of shared parameters and for all constructs using *RMSE*_*All*_ (i.e., fit *k*_*i*_ for each input *τ*_*B*_ and *τ*_*D*_ combinations for hairpin-series constructs). Then we minimize the sum of *RMSE*_*All*_ to find a global solution of shared input parameters. The thresholds in the *CI* calculations were set to the number of constructs to be used to fit each parameter.

#### Image analysis of smFISH-IF data

We developed a custom Matlab code (uLocalize) to perform smFISH-IF image analysis to identify translation sites intensity for mRNAs. Briefly, the mRNA channel is filtered with Laplacian of Gaussian filter to remove noise. The filtered image is thresholded to identify candidate spots, which are further quantified using Gaussian fitting to identify the location and intensity. The location of the RNA spot is used in the protein channel to draw a small region of interest (ROI) for potential TLS. To analyze the mature single proteins, the ROI containing potential translation sites were excluded. The protein channel is analyzed similarly as in the RNA channel: the positions and the integrated intensities of single proteins are identified and determined through Gaussian Fitting. To calculate the integrated intensity of TLS, the corresponding ROI is fitted with 3D Gaussian. To exclude mis-identified fitting to background signal, we imposed a minimal threshold of amplitude and maximal distance threshold to the corresponding mRNA. Finally, the integrated intensity of TLS is normalized to the single protein intensity in the same cell to calculate the number of nascent peptides. The program can be found on the Wu Lab Github site (https://github.com/binwulab/uLocalize.git).

#### Estimating number of nascent peptides from smFISH measurements

In our smFISH-IF measurement, we ignored mRNA-only populations as some of them could be impaired in terms of translational activity. Also the measured *FI*s may not be directly proportional to the number of ribosomes especially when the ribosomes locate at the beginning of mRNA due to the unsaturation of SunTags per nascent peptide, because we normalized the *FI*s with the mean *FI* value of the mature single nascent peptides (i.e., 24xSunTags in our SINAPs constructs). These two factors cause an inconsistency between our measured number of nascent peptides (#*NAPs*) and actual number of ribosomes (#*R*).

For the precise analysis of our smFISH-IF measurement, we need to consider the probability of having at least one #*NAPs* on the mRNA, as well as the correction of unsaturated nascent peptides. First, we considered the probability distribution of #*R, PDF*(#*R*), under the assumption of stochastic initiation and elongation events, to estimate the portion of mRNAs with no ribosome on them.

Let’s consider that a ribosome starts synthesizing a protein encoded in *L* codons for the constitutive initiation case. Under the stochastic elongation process with the elongation rate of *k*_*e*_ codons/s, the time required for the complete translation, *τ*_*t*_, follows a gamma distribution.

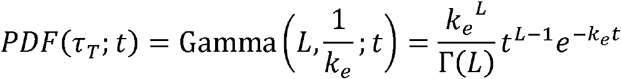

If *n* ribosomes were further initiated during *τ*_*t*_, the mRNA becomes having *N*-ribosomes on it after the first ribosome leaves. The probability of *N*-ribosome initiation (*I*_*N*_) follows the Poisson If ribosomes were further initiated during, the mRNA becomes having -ribosomes on it distribution with a mean value of *k*_*i*_ × *τ*_*t*_.

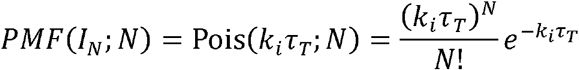

Thus, *PDF*(#*R*;*N*) can be derived by integrating all possible *τ*_*t*_ with the associated *I*_*N*_.

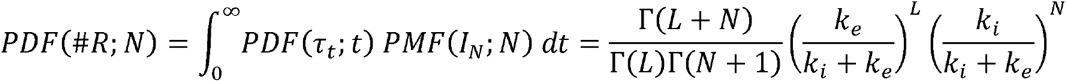

The expectation value of this *PDF*(#*R*;*N*) is resulted in the well-known relationship.

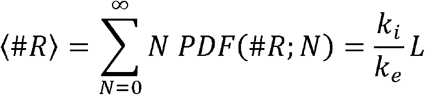

The probability of zero ribosome event, *P*(*N* = 0), at given, *k*_*i*_, *k*_*e*_ and *L* can be expressed as below, and further used to demonstrate the behavior of *P*(*N* = 0) as a function of *L* (Figure S3C).

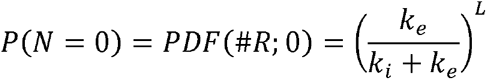

The number of synthesized SunTags (#*ST*), thus the *FI*, depends on the position of a ribosome on the mRNA. If a SunTag consists of *p*-codons and repeated for *q*-times at the beginning of ORF, #*ST* can be described as below.

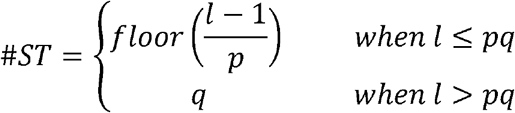

Here *l* is the position of a ribosome in terms of the codon number, and *floor* returns the nearest integer less than or equal to the input element. Mean number of SunTags per ribosome, <#*ST*>, can be derived by integrating all possible codon positions, with an assumption that the elongation rate is a constant through all codons (i.e., each codon has same probability having the ribosome).

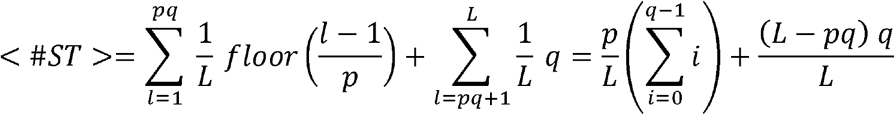

smFISH-IF analysis assumed that all ribosomes have *q*-SunTags, when we directly divide the *FI*s from TLS by mean *FI* of single nascent peptides to convert *FI* to #*NAPs* Thus the reported #*NAPs* are underestimating #*R* by <#*ST* >/*q*, which is equal to 0.6433 for our standard ORF length *L* = 841 codons.

In summary, our reported #*NAPs* are different from the actual number of ribosomes, can further converted to it by using above theoretical consideration. Figure S2C shows the relationship between #*NAPs* and #*R* with the considerations on both removing mRNA-only population and unsaturated nascent peptides for our standard ORF length (841 codons).

#### Analysis of RT-qPCR data

To generate standard curves for each qPCR target, the dilution factor was plotted against the Ct values. These data were fit by an exponential curve which was used to convert measured Ct values to a relative mRNA abundance. The triplicate-averaged NLuc reporter mRNA level for each sample was then normalized to the triplicate-averaged FLuc normalizing plasmid mRNA level and these ratios for each of two duplicate samples per condition were averaged to give a measure of the reporter mRNA level within each condition. All RT-qPCR and luciferase experiments were performed three independent times. In all cases for qPCR, the no-RT control reactions had Ct values which were >10 cycles higher than the corresponding RT samples, confirming little-to-no leftover genomic DNA.

#### Analysis of luciferase data

The ratio of NLuc:FLuc was measured for six technical replicate wells for each condition. The NLuc:FLuc ratio was then normalized to the mRNA ratio calculated above and these values for each hairpin insert were then normalized to the sextuplicate-averaged value for the no-insert condition to provide a measure of luciferase protein relative to RNA for each hairpin insert relative to the no-insert control.

#### Analysis of Phos-tag data

4E-BP and S6K phosphorylation was quantified using ImageJ (NIH). Band densities corresponding to phosphorylated and non-phosphorylated 4E-BP (*F*_*phoS*4*EBP*_) and S6K were quantified using profile plots to determine the fraction of each phosphorylated protein.

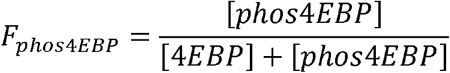

